# Refining sequence-to-expression modelling with chromatin accessibility

**DOI:** 10.1101/2025.02.11.637651

**Authors:** Orsolya Lapohos, Gregory J. Fonseca, Amin Emad

## Abstract

**Motivation:** Sequence-to-expression models typically do not consider chromatin accessibility, a major factor limiting gene regulation. We hypothesized that supplying accessibility as an input feature would allow a sequenceto-expression model to focus on important open regions of the genome.

**Results:** We found that the performance of such an augmented model was significantly better than that of sequence-only or accessibility-only models with similar architectures. Specifically, its ability to predict the expression of highly variable genes and gene expression in other cell types improved, and higher attribution scores in the input DNA sequences of the augmented model conformed to accessibility, enabling the learning of cell type-specific sequence patterns. Additionally, we show that fine-tuning a pre-trained sequence-only model with both sequence and accessibility can boost performance further and highlight the importance of sequencing depth in sequence-to-expression prediction.

**Availability and Implementation:** Source code is available on GitHub at https://github.com/lapohosorsolya/accessible_seq2exp.

## 1 Introduction

Metazoan gene expression is a complex process that is temporally and spatially orchestrated by a multitude of factors. At the lowest level, the blueprint for gene expression is ciphered into non-coding regulatory regions, defining the universe of DNA-binding proteins that can direct the regulation of each gene [1]. Although distal non-coding regions such as enhancers play important roles in controlling gene expression [2, 3], the promoter is arguably the master regulator. At the next level, the set of possible regulatory interactions is limited by context—chromatin accessibility and the set of available DNA-binding proteins both depend on cell type and environment [4]. Available regulatory proteins will compete and cooperate to bind accessible regions and ultimately activate or pause transcription initiation [5, 6]. Gene expression is further regulated during transcription and translation [1].

Despite the myriad of interactions regulating gene expression, sequence-to-expression models have been successful in predicting transcriptional activity from only promoter sequences [7, 8, 9, 10, 11, 12]. The 1D convolutional neural network (CNN) is naturally suited to the task of learning local sequence patterns and is the most frequently used base architecture for sequence-to-expression modelling. Recently, attention mechanism was also leveraged to capture long-range interactions in Enformer, a state-of-the-art model that predicts several genome tracks from long input sequences [13]. However, existing models do not make use of rate-limiting gene-regulatory factors, such as chromatin accessibility, in order to predict gene expression. Assay for transposase-accessible chromatin sequencing (ATAC-seq) detects open chromatin [14] and this data is increasingly available at single-cell resolution in parallel with RNAseq [15], providing both cell type-specific gene expression and regulatory context. Thus, we hypothesized that providing ATAC-seq as an additional input feature would focus a sequence-to-expression model on open regions of chromatin and thereby improve prediction of gene expression and *post hoc* model explanations.

While this idea is theoretically sound, we questioned whether a neural network could, in practice, learn sequence patterns alongside a strong predictor of gene expression such as chromatin accessibility. Consequently, the goal of this study is *not* to design a new model, but to take a step back and answer a specific set of questions pertaining to the proposed model augmentation strategy, which is compatible with virtually all sequence-to-expression architectures. Accordingly, we designed a proof-of-concept study to thoroughly eval-uate the effect of augmenting a vanilla sequence-to-expression model using chromatin accessibility as an input feature. For computational tractability, we selected a model with relatively few parameters: we opted to use a CNN-based architecture similar to Xpresso [9], rather than a transformer-based model, and to limit input sequence lengths to the 2kb regions flanking the transcription start sites of protein-coding genes. By doing so, we were able to train accessibility-augmented and ablated architectures on 12 different cell types from three independent human multiome RNAand ATACseq datasets with nested cross-validation, and perform additional scrambling experiments.

The contributions of this study are as follows. We showed that our accessibility-augmented model predicted the expression of held-out genes more accurately than models with similar architectures that only used sequence information or accessibility. This strategy also resulted in a greater improvement on the difficult task of highly variable gene expression prediction, while being able to generalize to other cell types. Additionally, models that used scrambled inputs performed significantly worse compared to our augmented model, revealing that all input data modalities contribute to the prediction performance. Importantly, we also showed that the magnitude of attribution scores obtained via the Shapley Additive Explanations DeepExplainer [16] in all DNA input channels increased in regions where the underlying chromatin was accessible, compared to the unfocused scores obtained for the sequence-only model. By analyzing *k*-mer attribution scores, we additionally showed that the augmented model was less reliant on CpG content, and that cell type-specific sequence patterns were better modelled when ATAC-seq was included. Finally, we demonstrated the importance of sequencing depth in the context of model performance and showed that a pre-training strategy could further boost performance. The strategies tested and proposed here can be adapted to other sequence-to-expression models with ease, potentially benefiting various downstream tasks.

## 2 Methods

### 2.1 Promoter sequences

The genome sequence FASTA and comprehensive gene annotation GTF files for the GRCh38.p14 reference genome (release 44) were downloaded from GENCODE [17]. Then, the annotation file was filtered to obtain the coordinates of “Ensembl canonical” transcripts for protein-coding genes, which were used to extract the coding strand sequence 1kb upstream and 1kb downstream of the start of the first exon (transcription start site, or TSS).

### 2.2 Datasets

Human single-nucleus multiome ATAC and gene expression datasets for peripheral blood mononuclear cells (PBMC), brain, and jejunum, as well as a human PBMC single-cell gene expression dataset with cell surface marker antibody-derived tags (ADTs) were sourced from 10x Genomics (Data availability). Dataset details are listed in Supplementary Table S1. Preprocessing and annotation of multiome and single-cell datasets are described in Supplementary Notes 1 and 2, respectively. Four major cell types were used from each dataset, shown in Supplementary Figure S1A-C. Cell type marker genes and numbers are indicated in Supplementary Tables S2 and S3, respectively.

### 2.3 Representing chromatin accessibility and gene expression for sequence-to-expression modelling

To obtain pooled cell type-specific ATAC tracks for each 2kb input sequence around the TSS, raw ATAC-seq fragments mapped to the cell type of interest were aligned to input sequence coordinates. The pooled cell typespecific ATAC tracks were smoothed using the 1D Gaussian filter function from the SciPy package (with standard deviation 20) [18] since the raw tracks were noisy, and Gaussian smoothing yielded appropriate scale-space representations for the task at hand. Track values were then min-max normalized by cell type. For other analyses, a summary metric termed auATAC was also defined to represent the accessibility of a gene promoter as the area under its normalized 2kb ATAC track.

To obtain a measure of gene expression by cell type, the proportion of cells of the cell type of interest with a nonzero unique molecular identifier (UMI) count was calculated for each gene. This metric, termed “GEx” in this study, represents the probability that a gene is expressed while ensuring that the output values range between 0 and 1. We opted to use gene expression probabilities rather than transcript abundance for several reasons. First, a probabilistic representation may be less biased by enhancer-mediated and post-transcriptional regulation of mRNA levels. Second, probabilities provide a more comparable measure for evaluation metrics and for cross-cell type prediction. Lastly, additive attribution scores obtained in *post hoc* analyses of trained models have interpretable meanings comparable between genes when the model output is a probability.

To analyze the similarity of gene expression or accessibility between cell types, the Pearson correlation coefficient between all GEx or auATAC values was calculated (Supplementary Figure S1D-F). Spearman correlation was used to determine concordance between GEx and auATAC (Supplementary Figure S1G-I).

In one of our analyses, we categorized genes based on gene expression and chromatin accessibility. The details of this method are described in Supplementary Note 3.

### 2.4 Model architecture

Figure 1 shows an overview of the model. One-hot-encoded DNA sequences with pooled and normalized ATAC-seq tracks serve as the 5 input channels to the neural network. The model outputs a single value that represents the expression of the input gene. The architecture of the neural network is based on Xpresso [9], but with a shorter input sequence (2kb instead of 10kb), an additional input channel, fixed depth and layer widths, and an additional dense layer (details in Supplementary Table S4 and Supplementary Figure S2). Our model was constructed with PyTorch v2.2 [19].

**Figure 1:**
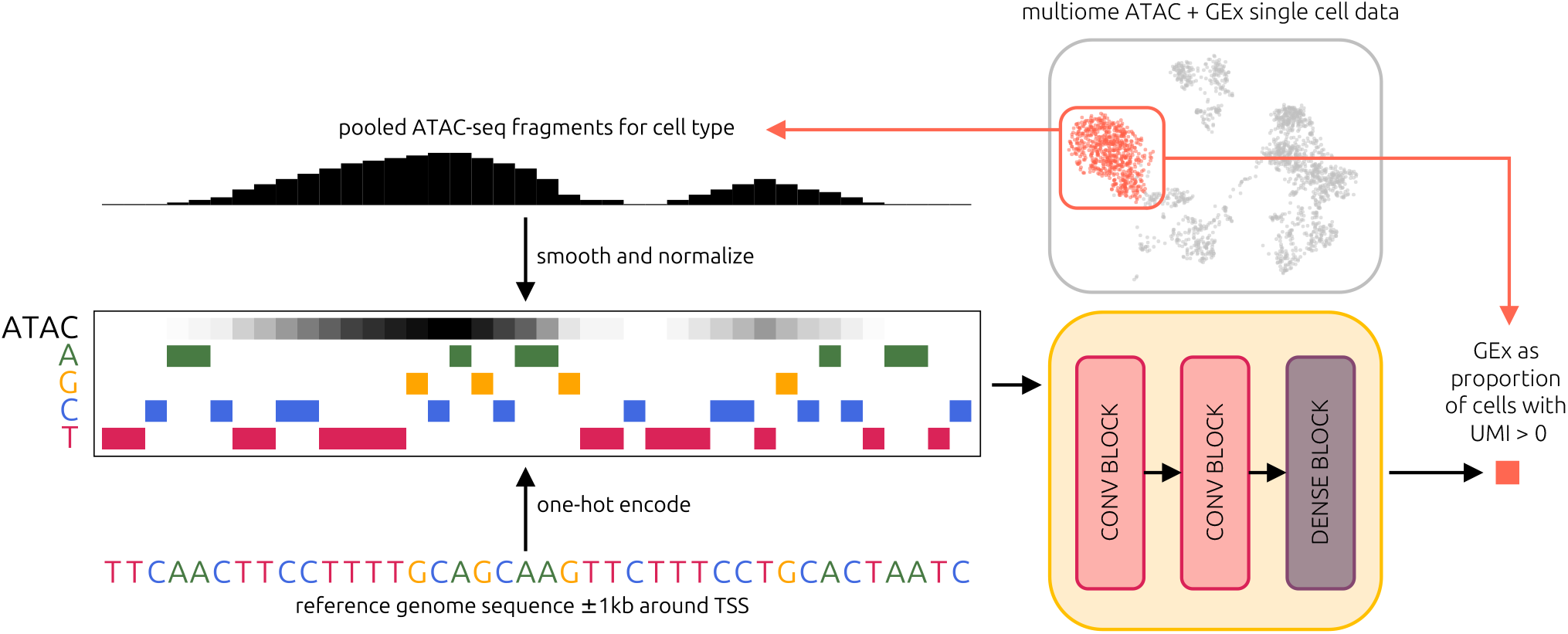
Overview of the ATAC-augmented sequence-to-expression model. Each input example consists of 5 vectors of length 2000, representing the promoter of a gene in a specific cell type. A pooled ATAC-seq track accompanies the one-hot encoded DNA nucleotides of the promoter sequence. For each input example, the model outputs a single value that represents the proportion of cells expressing the input gene.

For ablation experiments, model architecture remained unchanged, other than the input channels, which could consist of the cell type-specific ATAC track (1 channel), the one-hot-encoded DNA sequence of the promoter (4 channels), or both (5 channels).

### 2.5 Model training

Each model was trained and tested using 5-fold crossvalidation (CV) on the entire set of human autosomal protein-coding genes (19,095), where each test set consisted of unique genes. Training for each fold was repeated 5 times, each with different random seeds used for model initialization and splitting inner training and validation sets. Gene order was preserved across all datasets for each seed in each CV fold. The model was trained with mean squared error (MSE) loss, defined as

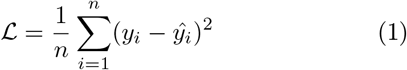

where *n* is the number of genes, *y*_*i*_ is the ground truth GEx of gene *i*, and *ŷ*_*i*_ is the predicted GEx for gene *i*. Training was guided by the Adam optimizer [20], with learning rate 5 × 10^−5^ and weight decay set to 10^−3^. Batch size was set to 512, and the training procedure was run for 500 epochs. Validation loss was computed every 4 epochs, and the model with the lowest validation loss was saved.

### 2.6 Fine-tuning of pre-trained models

An additional model training strategy involved a twostep procedure. First, a sequence-to-expression model was trained on sequence only, as described in the previous subsection. Then, these optimized weights were used to initialize a new full model, with input ATAC channel weights initialized randomly. This full model was then fine-tuned by training on both DNA and ATAC features. The fine-tuning procedure was performed for a maximum of 200 epochs, with reduced weight decay (10^−4^), a lower learning rate (10^−5^), and using the same 5-fold CV and random seeds. The model with the lowest validation loss was saved.

### 2.7 Evaluation

For each experiment, performance was assessed across 5 folds. For each fold, the average prediction across 5 models trained with different random seeds was evaluated. Reported means and standard deviations were calculated across the 5 folds. Four different metrics were calculated: Pearson correlation coefficient, Spearman correlation coefficient, MSE, and the coefficient of determination (*R*^2^).

### 2.8 Naïve prediction and evaluation

For each input gene promoter, the naïve predictor outputs its auATAC. Evaluation was performed using the test sets from the same 5-fold CV, calculating the Spearman correlation coefficient between the prediction (auATAC) and the ground truth GEx.

### 2.9 *Post hoc* model explanations

We obtained attribution scores for each held-out test sample using the SHapley Additive exPlanations (SHAP) DeepExplainer [16]. As background for each sample, we supplied 100 shuffled versions that preserved its dinucleotide frequencies, using the dinucleotide shuffler from the DeepLIFT package [21]. For background samples used to explain the DNA+ATAC model, values in the ATAC input channel were randomly shuffled as well. To summarize positional attribution scores by input channel, the mean SHAP score for each sequence position in each channel was calculated across 5 random seeds, for each CV fold. Then, means from all held-out test genes of the 5-fold CV were concatenated. Finally, the mean SHAP score was calculated at each position across all held-out test genes, for each input channel.

### 2.10 *k*-mer attribution analysis

We obtained attribution scores for *k*-mers with *k* = 2 and 6, where the score of a single *k*-mer instance in an input sequence is the mean SHAP score across the corresponding bases at each position. For every possible *k*-mer, we obtained scores at all instances within the input sequences, for all 25 trained models (5 CV folds and 5 random seeds), for one cell type. For *k* = 2, we took the mean attribution score across all instances and averaged these again across cell types to obtain an overall cell type-agnostic score for each dinucleotide. Then, we ranked 2-mers using these scores to compare model types. To analyze longer sequence patterns that may confer some cell type-specificity, we looked at 6-mers. For each 6-mer, we took the average across all its instances with positive SHAP scores to determine its role in positive regulation of gene expression in isolation.

Then, we ranked all 6-mers for each cell type and compared the top 10% pairwise using the Jaccard index, defined as |*A* ∩*B*| */*| *A* ∪*B*| for two sets *A* and *B*. We also repeated this analysis using all instances, regardless of score sign.

### 2.11 Statistical testing

The sets of held-out test genes used for evaluation across all 5 folds were preserved across all models. Thus, we used the one-sided Wilcoxon signed-rank test (from SciPy [18]) to test for statistical significance, when comparing performance of different models.

## 3 Results

### 3.1 ATAC-seq refined sequence-to-expression prediction

In order to determine whether providing chromatin accessibility as an additional input feature could improve sequence-to-expression modelling, we augmented a CNN-based model architecture similar to Xpresso [9] by including an extra input channel that represents a pooled ATAC-seq track for a specific cell type, in addition to the one-hot encoding of the input sequence (Figure 1). This architecture could be ablated easily by modifying only the set of input channels. We trained such models on sequence (DNA-only), accessibility (ATAC-only), and both (DNA+ATAC), to predict the proportion of cells within a cell type that express a gene (GEx).

First, we focused on DNA-only as our baseline sequence-to-expression model. This model achieved an average Pearson correlation of 0.3658 across the four major cell types in the PBMC dataset (Table 1, DNA only). Spearman correlations were higher for all four cell types, with an average of 0.5319. Performance was better on the brain and jejunum datasets overall, with average Pearson correlations of 0.4966 and 0.4573, and Spearman correlations of 0.5970 and 0.5620, respectively (Supplementary Table S5). The same trend was observed with MSE and *R*^2^ metrics.

**Table 1:**
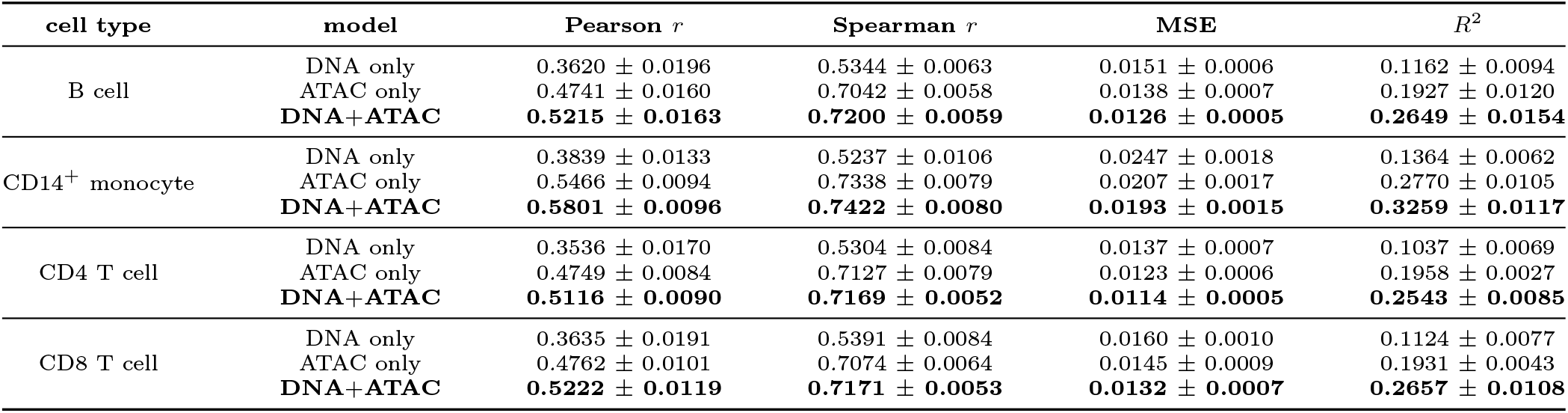
Results of ablation experiments with the PBMC dataset. Performance metrics on the held-out test sets are shown. MSE: mean squared error. All comparisons within cell type are significantly different by one-sided Wilcoxon signed-rank test (*p <* 0.05), except ATAC only versus DNA+ATAC Spearman *r* for CD4 T cell.

As expected, ATAC alone was a better predictor of gene expression than sequence alone (Table 1, ATAC only). This was already apparent in a naïve predictor that predicts GEx based on its auATAC (Methods). The average Spearman correlation of the naïve predictions with ground truth GEx for PBMCs was 0.694 (Supplementary Figure S3). The ATAC-only model achieved an average Pearson correlation of 0.4930 and Spearman correlation of 0.7145 on the PBMC dataset, improving on the naïve predictor.

Combining DNA with ATAC significantly and consistently improved model performance across all cell types (Table 1, DNA+ATAC; Supplementary Table S5; *p <* 0.05 based on one-sided Wilcoxon signed-rank tests, comparing DNA+ATAC to ablated models). The average Pearson and Spearman correlations for PBMCs were 0.5339 and 0.7241, for the brain dataset, they were 0.6621 and 0.7579, and for the jejunum dataset 0.5522 and 0.6925.

Since ATAC alone is a relatively strong predictor of GEx, we performed scrambling experiments to determine the contribution of DNA sequence to the DNA+ATAC model output. We scrambled one or both input features in the training sets with respect to the target genes, trained new models, and evaluated their performance on the unscrambled test sets. We also reevaluated predictions of models trained on unscrambled training sets using scrambled test sets as ground truth. In all scrambling experiments, performance was significantly worse than that of the unscrambled DNA+ATAC model (Figure 2; *p <* 0.05 based on one-sided Wilcoxon signed-rank tests).

**Figure 2:**
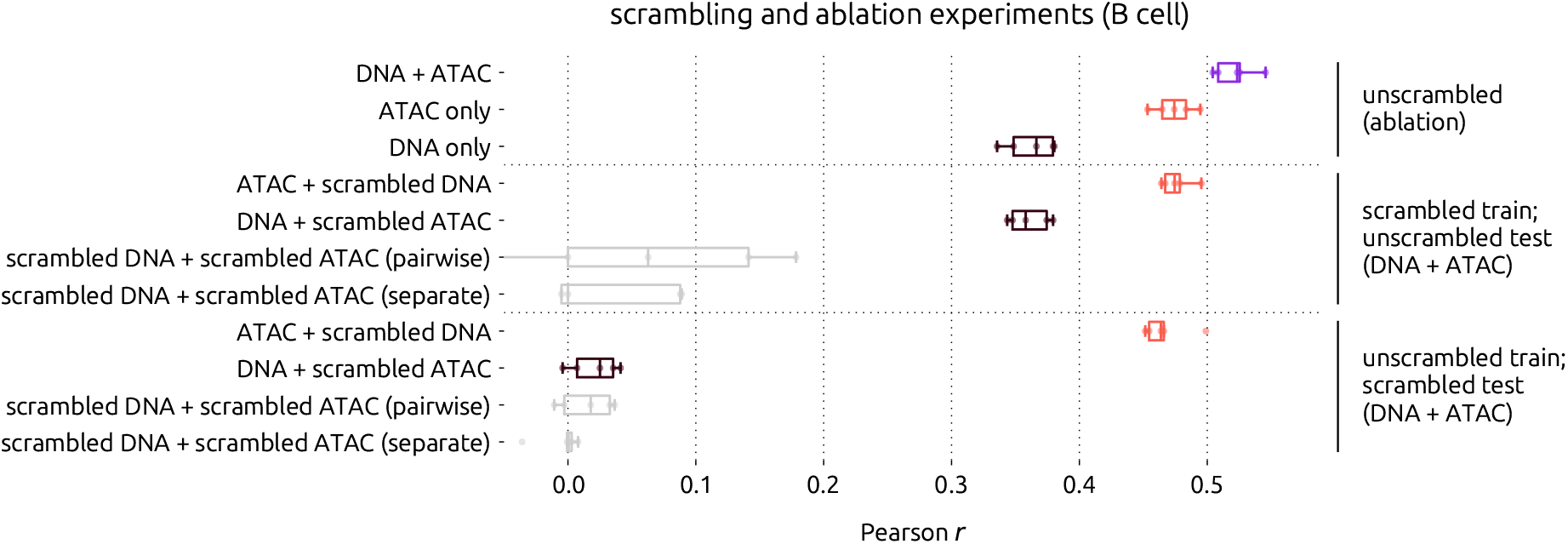
Scrambling experiments with B cell models. Pearson correlations of predicted versus ground truth GEx across 5-fold CV for each indicated ablated or scrambled model. Pairwise scrambling means the promoter sequence and ATAC track were kept together. Separate scrambling means promoter sequences, and ATAC tracks were scrambled independently. All vertically adjacent pairs of boxplots, except for the bottom pair, are significantly different by one-sided Wilcoxon signed-rank test (*p <* 0.05).

When the DNA inputs in the training set were scrambled, the DNA+ATAC model achieved approximately the same performance on the unscrambled test set as the unscrambled ablated ATAC-only model (mean Pearson *r* ≈ 0.47), indicating that unscrambled DNA is required to achieve the full performance of the DNA+ATAC model. Likewise, when the ATAC inputs in the training set were scrambled, similar performance was achieved on the unscrambled test set as the ablated DNA-only model (mean Pearson *r* ≈ 0.36). We also experimented with scrambling both inputs for training, to determine whether combinatorial effects between sequence patterns and accessibility could be learned by the augmented model. When inputs were scrambled pairwise (i.e.: DNA and ATAC of the same gene promoter were kept together) the model performed poorly, with mean Pearson *r* = 0.0492. When inputs were scrambled separately (independently with respect to the target gene), performance was significantly worse (*p* = 0.0339, onesided Wilcoxon signed-rank test), with mean Pearson *r* = − 0.0317. This suggests that, despite training on scrambled data, some predictive patterns can be learned when DNA and ATAC are complementary. Our findings were corroborated by a similar experiment where only the test set was scrambled (Figure 2). Altogether, results of the ablation and scrambling experiments showed that the input DNA sequences contributed meaningfully to the DNA+ATAC model output.

### 3.2 ATAC-seq feature improved cross-cell type and highly variable gene expression prediction

We evaluated generalizability by testing each trained cell type-restricted model on other cell types within the same dataset. In all three datasets, Pearson correlations on the cross-cell type prediction tasks were similar to those obtained when predicting on the same cell type, for both the ablated and augmented models (Figure 3A). These results indicate that the ATAC-seq input refined sequence-to-expression prediction from a generalization perspective as well. However, it should be noted that ground truth GEx values correlate highly between different cell types. The Pearson correlation coefficients between ground truth GEx of different cell types within the same dataset ranged from 0.699 to 0.981 (Supplementary Figure S1D-F, top). Cross cell-type Pearson correlation of auATAC was higher, with values between 0.817 and 0.992 (Supplementary Figure S1D-F, bottom). With such similarities, it is not surprising that sequenceto-expression models can generalize between cell types.

**Figure 3:**
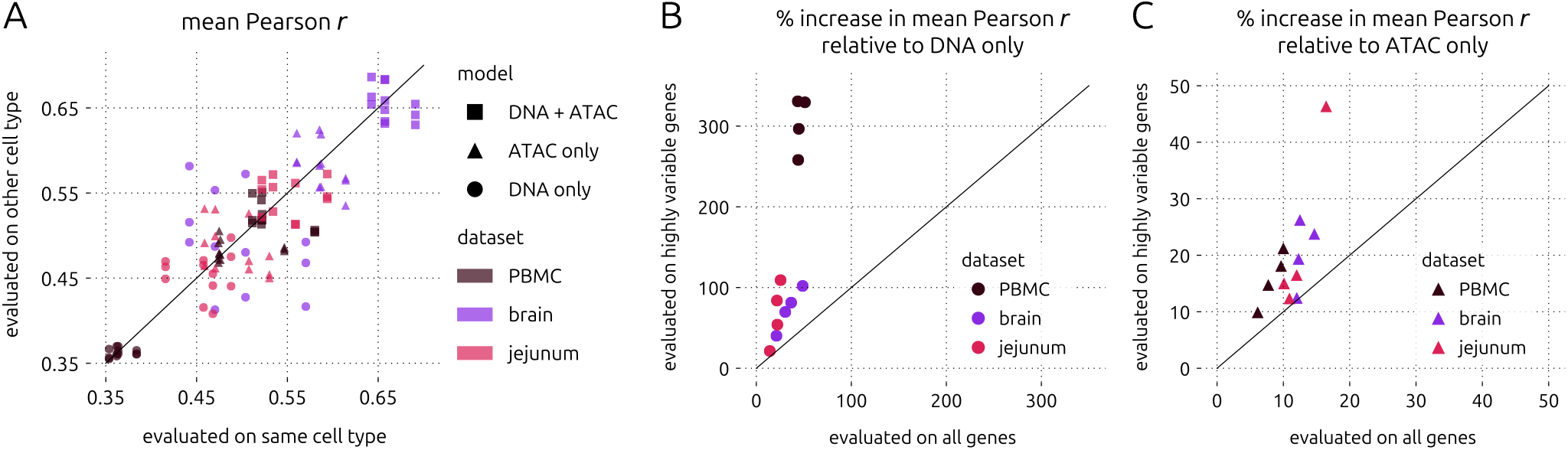
Cross-cell type and highly variable gene expression prediction results. A. Mean Pearson *r* of models evaluated on held-out test sequences using another cell type from the same dataset (tissue) versus on the cell type used for training. B. Percent increase in mean Pearson correlation from DNA only to DNA+ATAC for all major cell types in all three multiome datasets, evaluating on highly variable genes versus all genes. D. Same as C, but showing percent increase in mean Pearson correlation from ATAC only to DNA+ATAC. Solid diagonal lines indicate *y* = *x*.

In order to assess whether this observed generalizability is due to lowly variable (likely housekeeping) genes, we also evaluated the same models using subsets of the held-out test genes that were highly variable (Supplementary Note 1). As expected, all models had difficulty predicting the expression of highly variable genes, but models using ATAC were more resilient. When considering models trained and tested on the same cell type, the relative increase in mean Pearson correlation from DNA-only to DNA+ATAC was greater when evaluating on highly variable genes compared to all test genes (Figure 3B). We observed the same trend in the relative performance improvement from ATAC-only to DNA+ATAC (Figure 3C). Interestingly, we also observed these patterns on the cross-cell type prediction tasks (Supplementary Figures S4-7, Supplementary Data), although GEx of some cell types seemed to be inherently easier to predict. Taken together, these results suggest that the ATAC-augmented model is less biased by lowly variable genes while being capable of predicting on other cell types.

### 3.3 Positional attribution scores of input channels aligned with chromatin accessibility

To better understand our ATAC-augmented sequenceto-expression model, we examined attribution scores corresponding to each input as determined by the SHAP DeepExplainer [16], starting with the ATAC channel. Given that ATAC alone is a strong predictor of GEx— considering auATAC as a naïve predictor of GEx, as well as the performance of the ATAC-only model—we wanted to determine its contribution to model outputs for different categories of genes. We sorted genes into low, intermediate, and high bins, separately by GEx and auATAC (Figure 4A, Methods). Then, we intersected these bins to obtain 9 categories. For 97.9% of genes, GEx tended to increase with auATAC in CD4 T cells. A few genes (348) had high auATAC and low GEx, while a smaller number of genes had low auATAC and high GEx (60). We suspected that temporary silencing/pausing or long mRNA half-lives may explain these rare discrepancies. However, there was no concordance between an aggregated general mRNA halflife feature from a published meta-analysis [22] and the high GEx low ∩ auATAC category (Supplementary Figure S8A). Analyzing pausing indices calculated from a published PBMC CD4 T cell precision run-on sequencing (PRO-seq) dataset [23] (Supplementary Note 4) revealed that in the low GEx ∩ high auATAC category, only 25.2% of genes had a pausing index greater than 2 (Supplementary Figure S8B). Altogether, these observations suggest that neither mRNA half-life nor promoter-proximal pausing can explain all discrepancies between GEx and auATAC—these rare occurrences may be better explained by technical limitations.

**Figure 4:**
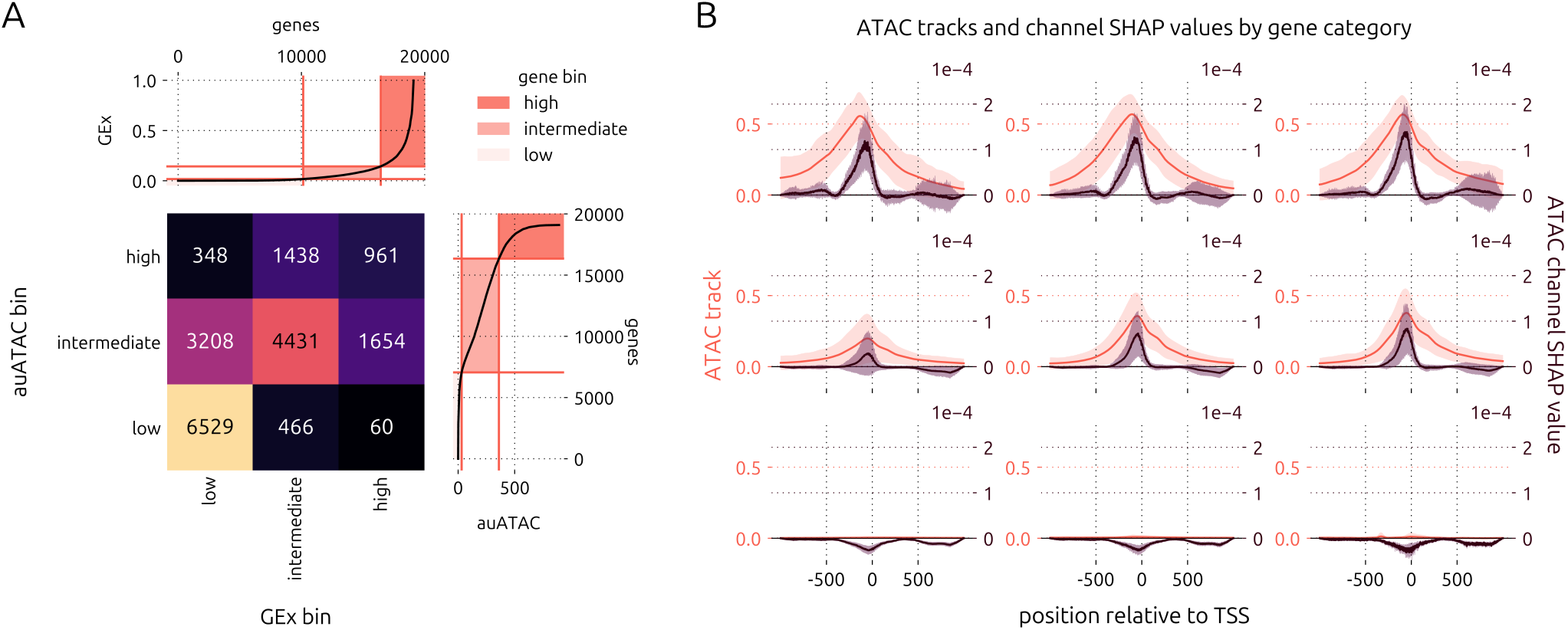
Chromatin accessibility and corresponding attribution scores in CD4 T cells, broken down by gene category. A. Gene contingency table generated by double knee point thresholding on GEx and area under ATAC (auATAC) for CD4 T cells. B. ATAC SHAP scores and ATAC input tracks for each category of the gene contingency table shown in A (DNA+ATAC model). Solid lines show means and shaded regions show standard deviations across all genes (after taking mean positional SHAP values across all random seeds).

Nevertheless, we computed positional attribution scores and averaged them for each gene category (Figure 4B). We observed that the ATAC input heavily influenced our model’s output, and that the largest mean ATAC attribution scores aligned with the mean ATAC track peak, slightly upstream of the TSS. Interestingly, relatively large values in the ATAC input track (~ 0.25) did not contribute positively to the model’s output in a small window (400-450bp) upstream of the TSS and in a wider window (100-400bp) downstream of the TSS. For genes with high auATAC, ATAC attribution scores were highly variable 500-1000bp downstream the TSS, indicating that accessibility in this section of the 5’ untranslated region was deemed relevant to the model. Overall, these results show that our DNA+ATAC model learned TSS-relative position-specific patterns in chromatin accessibility.

Next, we compared positional attribution scores in all input channels between the DNA-only and the DNA+ATAC models (Figure 5A). The SHAP scores of the DNA-only model were more distributed across the positions of the input sequence, while the DNA channels of the DNA+ATAC model had more variable SHAP scores that mirrored variability in the ATAC channel. We also calculated Spearman correlation coefficients of mean SHAP scores with the ATAC input track values at each position, across all test genes, for each channel (Figure 5B). Correlation coefficients were generally close to

**Figure 5:**
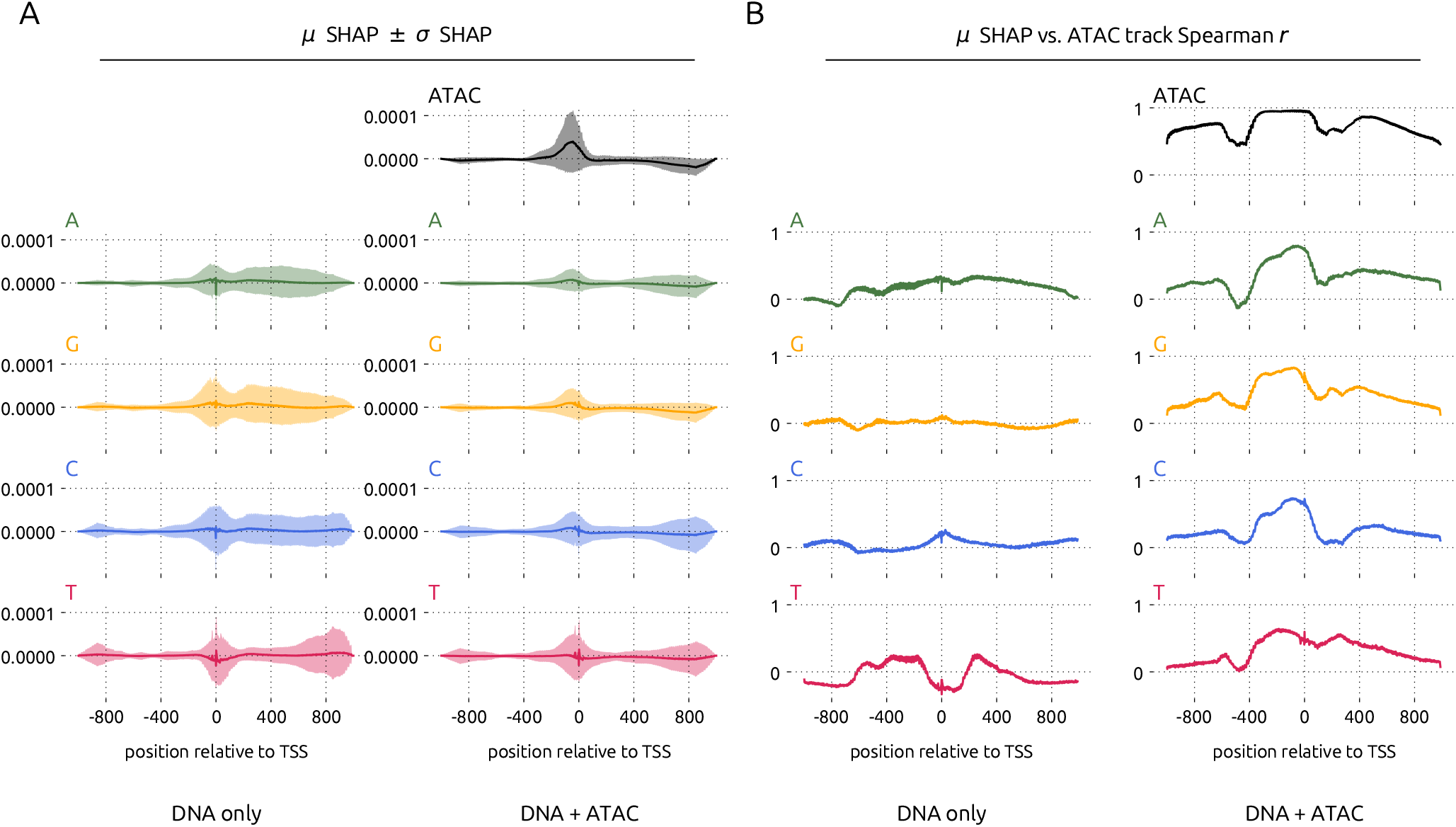
Positional attribution score distribution and correlation with accessibility across all input channels of the DNA-only and DNA+ATAC CD4 T cell models. A. *µ* and *σ* SHAP represent the mean and standard deviation of the positional SHAP scores in the indicated channel across 5 random seeds for all test genes (concatenated from all CV folds). Solid lines show *µ* and shaded regions indicate *σ*. B. *µ* SHAP vs. ATAC track Spearman *r* represents the positional Spearman correlation coefficient between *µ* SHAP and the input ATAC track across all test genes (concatenated from all CV folds).

zero in the DNA-only model, taking on values between − 0.461 and 0.372. In the DNA+ATAC model, SHAP scores in the ATAC channel correlated highly with the input ATAC track, with a minimum of 0.224 and maximum of 0.960, and coefficients in the DNA channels followed a similar pattern to those in the ATAC channel, with values between − 0.303 and 0.830. Hence, we surmise that the ATAC input feature guided the training of model weights in the DNA channels.

### 3.4 Replacing shallow single-nucleus GEx with deeper single-cell GEx improved performance

As the gene expression data obtained from multiome datasets is derived from single nuclei, we questioned whether model performance could be improved by replacing gene expression values with those from a matched more deeply sequenced single-cell dataset. When comparing the gene expression values between the multiome and single-cell RNA-seq PBMC datasets, we observed that the single-cell dataset was less sparse (Supplementary Figure S9A,B). GEx values derived from the single-cell dataset were also more highly correlated with auATAC values (Supplementary Figure S9C), than those from the multiome dataset (Supplementary Figure S1G). When we re-trained PBMC models on gene expression data from single-cells, we observed a significant improvement in the performance of the DNA+ATAC model, as well as the ablated models (Figure 6A, Supplementary Figure S10, Supplementary Data). These improvements also extended to highly variable gene expression prediction and generalizability across cell types. Further, we compared the correlation between positional attribution scores and chromatin accessibility obtained from models trained on CD4 T cells from the multiome and single-cell PBMC datasets (Supplementary Figure S11). While these correlational patterns were similar between the two datasets, the singlecell versions had more variability, which may be indicative of more nuanced attribution.

**Figure 6:**
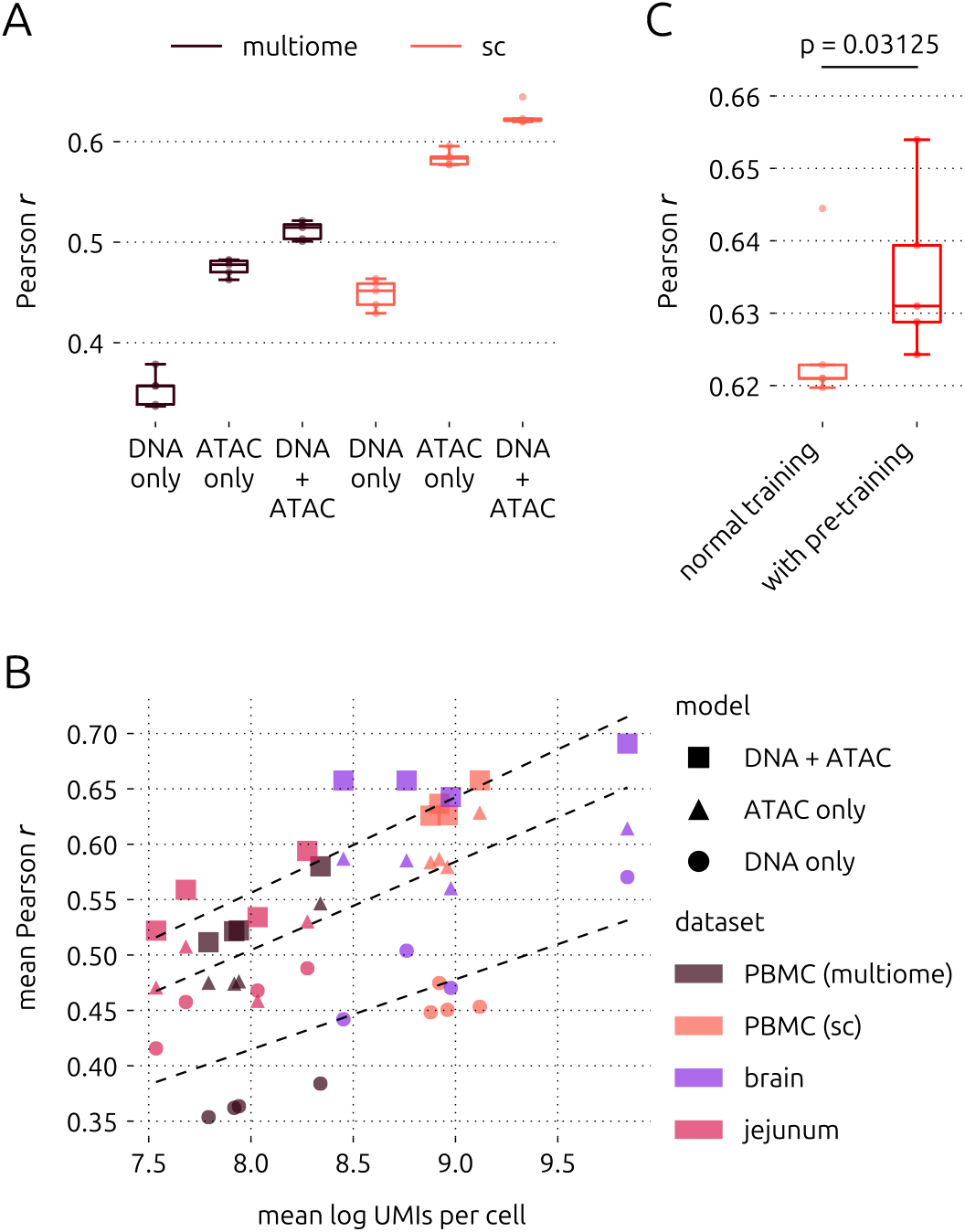
Performance improvement with single-cell GEx, sequencing depth, and pre-training. A. Performance improvement with single-cell versus multiome CD4 T cell data. Each comparison is significantly different (*p <* 0.05) by one-sided Wilcoxon signed-rank test. B. Relationship between model performance and mean log UMI counts. Each point represents the mean Pearson correlation achieved by a model trained and tested on one cell type, colour coded according to its dataset of origin. Square, triangle, and circle markers represent DNA+ATAC, ATAC only, and DNA only models, respectively. Dashed lines of best fit are shown for each model (top: DNA+ATAC, bottom: DNA only). C. Performance improvement with pre-training, using CD4 T cells (single-cell). “Normal training” refers to the DNA+ATAC model being trained starting from randomly initialized weights. “With pretraining” refers to the DNA+ATAC model being trained starting from weights copied from the DNA-only model. Boxplots show test results from 5-fold CV. Significance was determined by one-sided Wilcoxon signed-rank test.

### 3.5 UMI count affected model performance

After having trained sequence-to-expression models on several datasets, we suspected that differences in underlying statistics may have affected model performance. We investigated whether the number of reads per cell (Supplementary Figure S12), the number of pooled cells (Supplementary Table S3), or the variation in GEx values impacted model training. Plotting the mean Pearson correlation of each cell type-restricted model against the corresponding mean log UMIs per pooled cell, we observed that performance increased linearly with read depth (Figure 6B). This trend occurred in the DNA-only, ATAC-only, and DNA+ATAC models, with *R*^2^ values of 0.49, 0.79, and 0.82, respectively (Supplementary Figure S13, top). The number of pooled cells (Supplementary Figure S13, bottom left) did not seem to have an effect on performance, implying that the minimum number of pooled cells (185) was sufficient to obtain accurate gene expression probabilities. Finally, we found a trend between mean Pearson correlation and the variance-to-mean ratio (VMR) of GEx values for the ATAC-only and DNA+ATAC models (*R*^2^ = 0.68 and 0.53, respectively), but not the DNA-only model (Supplementary Figure S13, bottom right). This result further corroborates our finding that including accessibility as an input feature helps to explain some of the variability in gene expression. Altogether, these results demonstrate the importance of sequencing depth in sequence-to-expression modelling.

### 3.6 Pre-training strategy improved model performance

To explore additional strategies that could improve the learning of sequence patterns predictive of gene expression, we tested whether fine-tuning the pre-trained single-cell CD4 T cell DNA-only model with DNA and ATAC would help. We hypothesized that letting model weights reach an optimum based only on sequence prior to introducing the ATAC feature would improve performance. To test this hypothesis, we trained a new DNA+ATAC model, initializing it with weights copied from the corresponding DNA-only model (details in Methods). We found that there was indeed a statistically significant improvement in performance with the pre-training strategy, compared to the DNA+ATAC model trained from randomly intialized weights (Figure 6C; *p* = 0.0313, one-sided Wilcoxon signed-rank test). Additionally, we observed a slight change in the positional correlation of channel attribution scores with the input ATAC tracks (Supplementary Figure S11). With the pre-training strategy, Spearman correlation coefficients were generally similar to or lower than those obtained with the DNA+ATAC model. With the improvement in performance, these observations suggest that the pre-training strategy helps to balance information learned from sequence and accessibility. Thus, finetuning a pre-trained model with both sequence and accessibility is a viable strategy for boosting performance and it may allow the learning of more nuanced sequence patterns as determined by *post hoc* explainers.

### 3.7 Sequence pattern attribution improved with ATAC-seq input

Trained sequence-to-expression models are often used to infer the effects of short sequence patterns on gene expression. For an unbiased systematic comparison of sequence pattern attribution scores between our augmented DNA+ATAC and the DNA-only models, we analyzed all instances of *k*-mers in our input sequences. First, we ranked all 2-mers by their mean attribution scores and averaged them across PBMC cell types. We found many changes in the 2-mer importance score rankings between the two model types, the most noteworthy change being that predictions became less reliant on the presence of CpG dinucleotides in the augmented version (Fig. 7A). This finding is a good indicator that allowing the ATAC-seq input to explain some of the variability in gene expression enables the detection of less trivial sequence patterns. Interestingly, we also observed that reverse complements of dimers were often ranked very differently, indicating that *k*-mer orientation is relevant to gene expression prediction.

**Figure 7:**
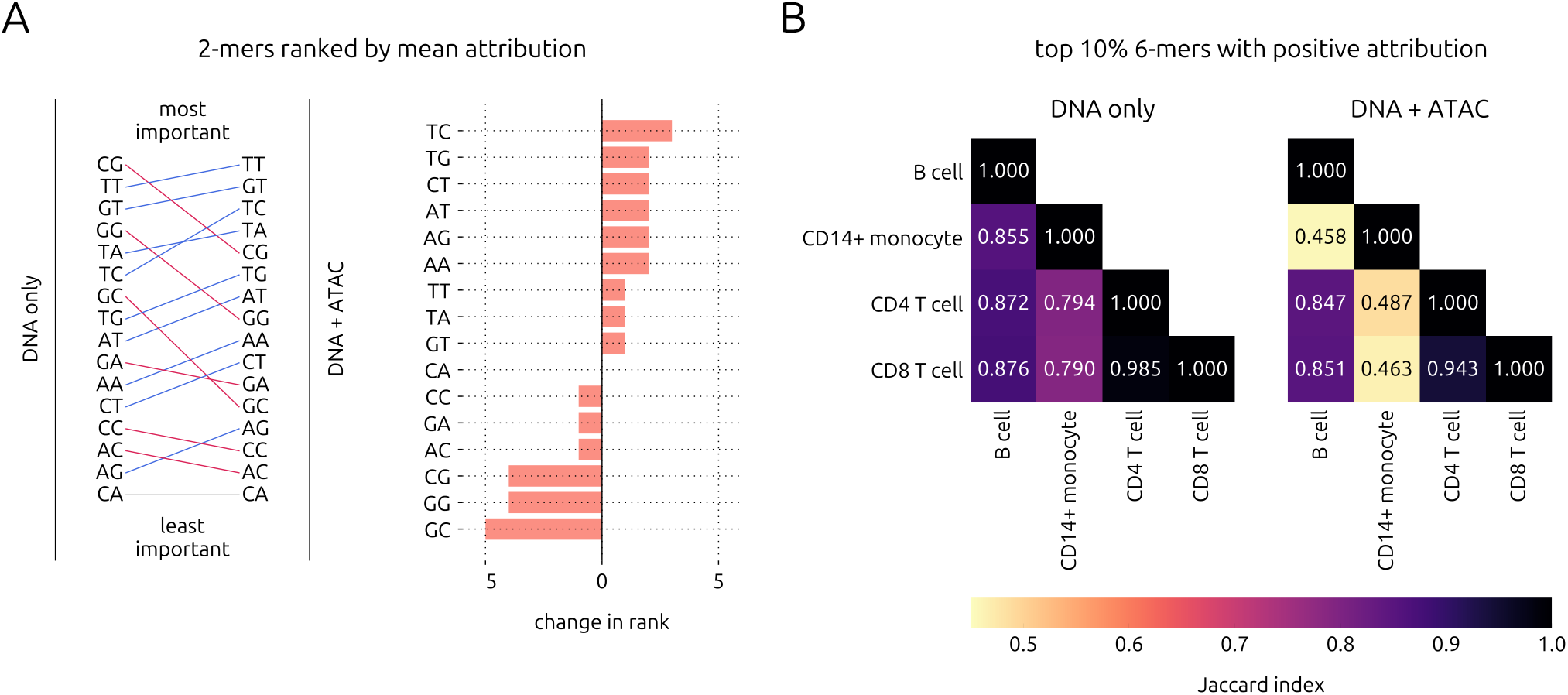
*k*-mer attribution comparison between the DNA-only and DNA+ATAC models trained on the PBMC (sc) dataset. A. Change in rank of mean 2-mer importance from the DNA-only to the DNA+ATAC models, across all folds, random seeds, and cell types. B. Jaccard index of the top 10% of 6-mers with positive attribution between the PBMC cell types.

Since cell type-specific gene expression is governed by higher-level patterns, we also compared 6-mer importances across cell types. We found that the overlaps between the top 10% of 6-mers among instances with positive attribution in the PBMC cell types approximated the biological similarities of these cell lineages more closely when we used the attribution scores from the DNA+ATAC models (Fig. 7B). The CD4 and CD8 T cell subsets are known to be highly similar while B cells and T cells are both members of the lymphoid lineage, sharing some transcriptional programs. CD14^+^ monocytes are members of the myeloid lineage, and are therefore the most distant from the three other cell types tested here. These relations are best reflected in the pairwise Jaccard indices between the top 6-mers obtained using the augmented models. We observed a similar pattern when repeating this analysis using mean scores across all 6-mer instances, including negative ones (Supplementary Figure S14). Altogether, our *k*-mer analyses demonstrate that sequence pattern attribution improved with the addition of ATAC-seq as an input feature.

## 4 Discussion

Sequence-to-expression models are not expected to capture all variation in gene expression on their own. Many layers of regulation restrict the realm of possibilities defined by a DNA sequence, and these layers cannot be known without measuring the cellular context to a sufficient degree. For instance, when Xpresso [9] and scXpresso [12] incorporated mRNA half-life features, gene expression prediction improved. Although general halflife features are not context-specific, they contribute an additional set of “rules” to define transcriptional dynamics. As this field moves towards building more complex models of gene expression, more variation will be explained by accounting for every possible regulatory layer, both sequence-inherent, and context-specific. Here, we demonstrated that augmenting input DNA sequences with chromatin accessibility captured in multiome single-cell datasets refined sequence-to-expression modelling. In all datasets we used, performance consistently and significantly improved not only with respect to the baseline DNA-only model, but also with respect to the ATAC-only model.

A major concern we had with the concept of feeding accessibility as an additional input rather than output to the model, was that accessibility alone may be too strong of a predictor of gene expression. As a result, it could be too easy for model weights to converge at a local minimum that lacks sequence-related knowledge during training. However, our concerns were alleviated after confirming, via ablation and scrambling experiments, that the input DNA sequences had contributed a significant amount of information. Moreover, the finding that the relative improvement from both DNA-only and ATAC-only to DNA+ATAC increased for highly variable genes (Figure 3C) implies that our strategy lessened biases by lowly variable (likely housekeeping) genes. This analysis also raises the question of whether sequence patterns in housekeeping gene promoters differ from those encoded in promoters of more specialized genes—future studies could further investigate this idea.

Most studies on sequence-based modelling have not reported generalizability results. A notable exception is a recent publication which introduced EpiBERT, a multi-modal transformer for cell type-agnostic predictions [24]. Here, we found that the performance of each model type on other cell types within the same dataset was similar to its performance on the same cell type. Further, these trends were not affected when evaluating on only highly variable genes. Thus, we suspect that either (1) the learned sequence patterns within accessible promoters were general, and/or (2) variability within tissue is too low to capture cell-type specific patterns. Follow-up studies are needed to further investigate these hypotheses. Nevertheless, improved modelling of highly variable genes was a positive outcome of our approach.

Taking our analysis further, attribution scores in the DNA input channels of the augmented sequence-to-expression model increased at accessible sequence positions. This result contrasted with the attribution scores for the DNA-only model which were more widely distributed across input sequence positions. We also demonstrated that adding ATAC-seq as an input feature reduced reliance on CpG content, a trivial sequence feature, and allowed for the detection of cell type-specific 6-mers despite looking no further than ±1kb from the TSS. Together, these results suggest that our augmentation strategy may result in improved inference of motif grammar and prediction of single nucleotide variant effects in future studies investigating larger models.

Lastly, we found two extra steps that improved model performance. The first of these steps involved the replacement of GEx values from relatively shallow singlenucleus multiome data with those from more deeply sequenced single-cell data. We also showed that model performance improves linearly with mean log UMIs per pooled cell, but it is worth mentioning that differences between single-nucleus and single-cell RNA-seq (see [25, 26]) may have considerable effects on ground truth and therefore model accuracy. The second step was the fine-tuning of a pre-trained DNA-only model using both DNA and ATAC inputs. We speculate that the reason for this improvement is that delaying the use of the strongly predictive accessibility feature might prevent distraction from learning subtle sequence patterns. However, we did not observe major changes in the positional correlation of channel attribution scores with the ATAC input track using the pre-training strategy, compared to normal training. Small decreases in correlation were observed in short segments of the promoter sequences, taking on a seemingly more balanced profile in between the DNA-only model and the DNA+ATAC model. This result implies that applying the fine-tuning approach to existing sequence-to-expression models could yield refined results with minimal effort.

Our study has the following limitations. Firstly, techniques that measure chromatin accessibility each have biases, so using alternative (or combined) accessibility data measured by DNase-seq may lead to different learned patterns [27]. Another limitation of our input data is that sequencing depth has a major impact on performance, and obtaining deeply sequenced single-cell data is costly. However, the concepts demonstrated in this study can be applied to bulk data as well, provided that both RNA-seq and ATAC-seq are available for a homogeneous biological sample. We also note that our input sequences are relatively short, and most sequenceto-expression models use much longer input sequences.

On the other hand, models using long input sequences have been shown to ignore effects of enhancers [28]. Since enhancers affect highly variable genes in particular [29], exploring new architectures and strategies capable of learning long-range effects is still important. Based on our findings, it is possible that augmenting a long-range model with ATAC-seq as an additional input feature could result in similar improvements that also take enhancers into account.

In conclusion, we added context to a plain sequence-to-expression model by including chromatin accessibility as an input feature. By modelling gene expression as a function of both sequence and accessibility, learning sequence patterns predictive of highly variable gene expression became easier and attribution scores of DNA input channels became aligned with accessibility. We are convinced that this portable sequence-to-expression augmentation strategy would benefit larger models and result in the inference of more nuanced cell type-specific regulatory grammar and improved single nucleotide variant effect prediction.

## 5 Data availability

Multiome human PBMC data were obtained from https://www.10xgenomics.com/resources/datasets/pbmc-from-a-healthy-donor-no-cell-sorting-10-k-1-standard-2-0-0. Multiome human brain data were obtained from https://www.10xgenomics.com/resources/datasets/frozen-human-healthy-brain-tissue-3-k-1-standard-2-0-0.

Multiome human jejunum data were obtained from https://www.10xgenomics.com/resources/datasets/human-jejunum-uclei-isolated-with-chromium-nuclei-isolation-kit-saltyez-protocol-and-10x-complex-tissue-dp-ct-sorted-and-ct-unsorted-1-standard.

The single-cell PBMC dataset with ADTs was obtained from https://www.10xgenomics.com/datasets/5k-human-pbmcs-stained-with-totalseq-B-human-TBNK-cocktail-NextGEM.

The aggregated mRNA half-life feature for human genes was obtained from a previously published meta-analysis [22] (file: 13059_2022_2811_MOESM3_ESM.xlsx).

The human PBMC CD4 T cell PRO-seq data were obtained from a previous publication by Danko et al. [23], downloaded from the Gene Expression Omnibus with accession number GSE85337.

## 6 Code availability

Implemented code is available on GitHub, at https://github.com/lapohosorsolya/accessible_seq2exp and also on Figshare, https://doi.org/10.6084/m9.figshare.28836278.

## 8 Supplementary material

Supplementary Notes, Tables and Figures in PDF format and Supplementary Data in Excel sheets are available online.

## 9 Author contributions statement

All authors contributed to the design of the study and the writing of the manuscript. A.E. and G.J.F. supervised the study. O.L. performed the experiments and analyzed the results. O.L., G.J.F., and A.E. read and approved the final manuscript.

