## Supplement for "Refining sequence-to-expression modelling with chromatin accessibility"

**Supplementary Tables** **4**

**Supplementary Figures** **10**

**References** **23**

### Supplementary Notes

#### 1 Processing and annotation of multiome data

Human single nucleus multiome ATAC and gene expression datasets for peripheral blood mononuclear cells (PBMC), brain, and jejunum were sourced from 10x Genomics (Supplementary Table S1, Data availability). For each multiome dataset, the files with suffixes `filtered_feature_bc_matrix.h5` and `atac_fragments.tsv` were downloaded (see Data availability).

In order to annotate nuclei by cell type, each gene expression matrix was analyzed with the Scanpy [Wolf et al., 2018] package. Briefly, cells with a proportion of mitochondrial unique molecular identifiers (UMIs) greater than 5% of total UMIs were removed, and multiplets were filtered using a total UMI threshold of two standard deviations above the mean. Then, gene expression values were library size-normalized and log-transformed for the identification of highly variable genes using default settings in Scanpy. Nuclei were clustered using the Leiden algorithm [Traag et al., 2019], and ranked differentially expressed genes were obtained between each cluster by Mann-Whitney test. Clusters were annotated manually, by comparing cluster-specific genes with cell type marker genes found by literature search and corroborated with The Human Protein Atlas [Karlsson et al., 2021, Uhlen et al., 2019]. Cell type marker genes are indicated in Supplementary Table S2. Four major cell types were used from each dataset. Among PBMCs, we chose B cells, $CD14^+$ monocytes, CD4 T cells, and CD8 T cells. Brain cells were divided into astrocytes/microglia, oligodendrocytes, neurons, and oligodendrocyte precursor cells (OPCs). Cells from the jejunum were divided into epithelial, stromal (immune), stromal (other), and intestinal stem cells (ISCs). Uniform Manifold Approximation and Projection (UMAP) [Mcinnes et al., 2020] graphs of gene expression in each multiome dataset are shown in Supplementary Figure S1A-C, with each major cell type indicated by colour. Cell numbers by cell type are indicated in Supplementary Table S3.

#### 2 Processing and annotation of single-cell data

A human PBMC single-cell gene expression dataset with cell surface marker antibody-derived tags (ADTs) was also obtained from 10x Genomics (Supplementary Table S1, Data availability). ADTs were denoised and scaled by background using the DSB method [Mulè et al., 2022] built into the MUON package [Bredikhin et al., 2022]. Cells were then filtered, transformed, and clustered based on gene expression, as described in the previous section. The levels of cell type-specific DSB-normalized ADTs in each cluster were then used to annotate the same four major cell types as in the PBMC multiome dataset (numbers in Supplementary Table S3).

#### 3 Gene binning method

We grouped genes into 9 categories based on gene expression and chromatin accessibility. For each dataset, we used an automated knee-point detection algorithm to bin genes by area under the normalized 2kb ATAC input track (auATAC) within the input sequence range and by GEx (probability of gene expression), separately. Briefly, genes were sorted in ascending order, for each feature. Then, the kneed package [Satopa et al., 2011] was used to normalize feature values and obtain a difference curve. The maximum of this difference curve was chosen as the first knee point. To obtain a second knee point, the sorted list of genes was truncated using the first knee point and the knee-point detection process was repeated. The two knee points calculated for each feature (auATAC and GEx) were used to sort genes into low, intermediate, and high bins. Finally, a gene contingency table was obtained by intersecting these bins between auATAC and GEx.

#### 4 Promoter-proximal pausing analysis

Three human PBMC CD4 T cell precision run-on sequencing (PRO-seq) samples published previously [Danko et al., 2018] were downloaded in the form of pre-processed BigWig files (see Data availability). Since these had been mapped to a different reference genome, corresponding gene coordinates were extracted from the GENCODE GRCh37 backmap reference GTF file [Frankish et al., 2022]. Then, the three samples were aggregated into one before calculating pausing indices. The pausing index for each gene was calculated by dividing the number of reads within the first 1kb downstream the TSS by the number of reads along the rest of the gene body (on the coding strand, and only for genes at least 3kb in length) as described elsewhere [Adelman and Lis, 2012].

### Supplementary Tables

S1 Dataset sequencing depth and pre-processing statistics . . . . . . . . . . . . . 5

S2 Cell type marker genes used to annotate major cell types . . . . . . . . . . . 6

S3 Major cell types identified and number pooled in each dataset . . . . . . . . 7

S4 Architectural details . . . . . . . . . . . . . . . . . . . . . . . . . . . . . . 8

S5 Ablation experiment results for all multiome datasets . . . . . . . . . . . . . 9

**Supplementary Table S1:** Sequencing depth and pre-processing statistics of datasets used in this study. ATAC: assay for transposase-accessible chromatin. GEX: gene expression. PBMC: peripheral blood mononuclear cells.

| Alias | 10x Genomics dataset | 10x Genomics pipeline version | Estimated cell number | ATAC mean raw read pairs per cell | ATAC median HQ fragments per cell | GEX mean raw read pairs per cell | GEX median UMI counts per cell | Year published |
|---|---|---|---|---|---|---|---|---|
| PBMC (multiome) | PBMC from a Healthy Donor - No Cell Sorting (10k) | v2.0.0 | 12,012 | 38,103 | 11,769 | 69,129 | 3,300 | 2021 |
| PBMC (sc) | 5k Human PBMCs Stained with TotalSeqTM-B Human TBNK Cocktail, Chromium NextGEM Single Cell 3’ | v8.0.0 | 5,078 | NA | NA | 71,818 | 7,612 | 2024 |
| brain | Flash-Frozen Human Healthy Brain Tissue (3k) | v2.0.0 | 3,233 | 95,897 | 22,881 | 122,335 | 6,966 | 2021 |
| jejunum | Human Jejunum Nuclei Isolated with Chromium Nuclei Isolation Kit, SaltyEZ Protocol, and 10x Complex Tissue DP (CT Sorted and CT Unsorted) | v2.0.2 | 10,640 | 58,571 | 8,340 | 17,307 | 3,676 | 2023 |

**Supplementary Table S2:** Cell type marker genes used to annotate major cell types in multiome datasets. PBMC: peripheral blood mononuclear cells. OPC: oligodendrocyte precursor cell. ISC: intestinal stem cell.

| Tissue | Major cell type | Relevant marker genes |
|---|---|---|
| PBMC | B cell | *CD19, MS4A1* |
| | $CD14^+$ monocyte | *CD36, CSF3R, S100A9, SLC8A1, VCAN* |
| | CD4 T cell | *CD4, IL7R, INPP4B, LEF1* |
| | CD8 T cell | *CD8A, IL7R, LEF1, NELL2* |
| brain | astrocyte/microglia | *AQP4, CD83, MERTK, P2RY12* |
| | oligodendrocyte | *PLP1* |
| | neuron | *CNR1, GAD2, SNAP25, SYNPR* |
| | OPC | *PDGFRA* |
| jejunum | epithelial | *BEST4, DEFA5, PTPRN2, SLC15A1, SLC12A2* |
| | stromal (immune) | *SEMA4D, MERTK* |
| | stromal (other) | *CDH19, LRMDA, MYH11, RSPO3, SYT1* |
| | ISC | *LRMDA, MET* |

**Supplementary Table S3:** Major cell types identified and number pooled in each dataset. PBMC: peripheral blood mononuclear cells. OPC: oligodendrocyte precursor cell. ISC: intestinal stem cell.

| Dataset | Identified major cell type | Cell number |
|---|---|---|
| PBMC (multiome) | B cell | 888 |
| | $CD14^+$ monocyte | 3295 |
| | CD4 T cell | 2225 |
| | CD8 T cell | 1144 |
| PBMC (single cell) | B cell | 272 |
| | $CD14^+$ monocyte | 1352 |
| | CD4 T cell | 1510 |
| | CD8 T cell | 551 |
| brain | astrocyte/microglia | 712 |
| | oligodendrocyte | 1757 |
| | neuron | 574 |
| | OPC | 185 |
| jejunum | epithelial | 5882 |
| | stromal (immune) | 1469 |
| | stromal (other) | 1059 |
| | ISC | 693 |

**Supplementary Table S4:** Architectural details. Layer types correspond to `torch.nn` module names. In the input layer, there are 1, 4, or 5 channels, depending on whether accessibility, sequence, or both are included. $p$ represents the probability of nodes to be zeroed out, $C_{\text{in}}$ represents the number of input channels, $C_{\text{out}}$ represents the number of output channels (filters), $k$ represents kernel size, $s$ represents stride, and $d$ represents dilation.

| Block | Layer Type | Parameters | Output Shape |
|---|---|---|---|
| | Input | | $x \times 2000 \mid x \in \{1, 4, 5\}$ |
| | Dropout | $p = 0.5$ | |
| 1 | Conv1d | $C_{\text{in}} = 5$, $C_{\text{out}} = 128$, $k = 6$, $s = 1$, $d = 1$ | $128 \times 1996$ |
| | MaxPool1d | $k = 8$, $s = 8$, $d = 1$ | $128 \times 249$ |
| 2 | Conv1d | $C_{\text{in}} = 128$, $C_{\text{out}} = 64$, $k = 9$, $s = 1$, $d = 2$ | $64 \times 233$ |
| | MaxPool1d | $k = 8$, $s = 8$, $d = 1$ | $64 \times 29$ |
| | Flatten | | 1856 |
| | Dropout | $p = 0.5$ | |
| 3 | Linear | | 1856 |
| | ReLU | | |
| | Dropout | $p = 0.5$ | |
| | Linear | | 64 |
| | ReLU | | |
| | Dropout | $p = 0.5$ | |
| | Linear | | 1 |

**Supplementary Table S5:** Ablation experiment results for all multiome datasets. Performance metrics on the held-out test sets are shown. MSE: mean squared error. All comparisons within cell type are significantly different by one-sided Wilcoxon signed rank test ($p < 0.05$), except ATAC only versus DNA+ATAC Spearman $r$ for CD4 T cell.

| Dataset | Cell type | Model | Pearson $r$ | Spearman $r$ | MSE | $R^2$ |
|---|---|---|---|---|---|---|
| PBMC | B cell | DNA only | 0.3620 ± 0.0196 | 0.5344 ± 0.0063 | 0.0151 ± 0.0006 | 0.1162 ± 0.0094 |
| | | ATAC only | 0.4741 ± 0.0160 | 0.7042 ± 0.0058 | 0.0138 ± 0.0007 | 0.1927 ± 0.0120 |
| | | **DNA+ATAC** | **0.5215 ± 0.0163** | **0.7200 ± 0.0059** | **0.0126 ± 0.0005** | **0.2649 ± 0.0154** |
| | $CD14^+$ monocyte | DNA only | 0.3839 ± 0.0133 | 0.5237 ± 0.0106 | 0.0247 ± 0.0018 | 0.1364 ± 0.0062 |
| | | ATAC only | 0.5466 ± 0.0094 | 0.7338 ± 0.0079 | 0.0207 ± 0.0017 | 0.2770 ± 0.0105 |
| | | **DNA+ATAC** | **0.5801 ± 0.0096** | **0.7422 ± 0.0080** | **0.0193 ± 0.0015** | **0.3259 ± 0.0117** |
| | CD4 T cell | DNA only | 0.3536 ± 0.0170 | 0.5304 ± 0.0084 | 0.0137 ± 0.0007 | 0.1037 ± 0.0069 |
| | | ATAC only | 0.4749 ± 0.0084 | 0.7127 ± 0.0079 | 0.0123 ± 0.0006 | 0.1958 ± 0.0027 |
| | | **DNA+ATAC** | **0.5116 ± 0.0090** | **0.7169 ± 0.0052** | **0.0114 ± 0.0005** | **0.2543 ± 0.0085** |
| | CD8 T cell | DNA only | 0.3635 ± 0.0191 | 0.5391 ± 0.0084 | 0.0160 ± 0.0010 | 0.1124 ± 0.0077 |
| | | ATAC only | 0.4762 ± 0.0101 | 0.7074 ± 0.0064 | 0.0145 ± 0.0009 | 0.1931 ± 0.0043 |
| | | **DNA+ATAC** | **0.5222 ± 0.0119** | **0.7171 ± 0.0053** | **0.0132 ± 0.0007** | **0.2657 ± 0.0108** |
| brain | astrocyte/microglia | DNA only | 0.5039 ± 0.0091 | 0.5933 ± 0.0079 | 0.0273 ± 0.0011 | 0.2241 ± 0.0104 |
| | | ATAC only | 0.5854 ± 0.0045 | 0.7096 ± 0.0031 | 0.0243 ± 0.0013 | 0.3086 ± 0.0089 |
| | | **DNA+ATAC** | **0.6574 ± 0.0077** | **0.7506 ± 0.0074** | **0.0203 ± 0.0010** | **0.4211 ± 0.0069** |
| | oligodendrocyte | DNA only | 0.4420 ± 0.0026 | 0.5895 ± 0.0071 | 0.0249 ± 0.0012 | 0.1722 ± 0.0079 |
| | | ATAC only | 0.5869 ± 0.0163 | 0.7432 ± 0.0046 | 0.0208 ± 0.0014 | 0.3087 ± 0.0174 |
| | | **DNA+ATAC** | **0.6575 ± 0.0141** | **0.7679 ± 0.0068** | **0.0173 ± 0.0013** | **0.4248 ± 0.0170** |
| | neuron | DNA only | 0.5703 ± 0.0097 | 0.6251 ± 0.0089 | 0.0529 ± 0.0006 | 0.2980 ± 0.0037 |
| | | ATAC only | 0.6142 ± 0.0060 | 0.6966 ± 0.0040 | 0.0520 ± 0.0013 | 0.3106 ± 0.0106 |
| | | **DNA+ATAC** | **0.6909 ± 0.0045** | **0.7520 ± 0.0039** | **0.0401 ± 0.0007** | **0.4683 ± 0.0056** |
| | OPC | DNA only | 0.4703 ± 0.0153 | 0.5802 ± 0.0123 | 0.0361 ± 0.0005 | 0.1923 ± 0.0098 |
| | | ATAC only | 0.5603 ± 0.0103 | 0.7168 ± 0.0043 | 0.0320 ± 0.0009 | 0.2846 ± 0.0100 |
| | | **DNA+ATAC** | **0.6426 ± 0.0109** | **0.7612 ± 0.0019** | **0.0267 ± 0.0009** | **0.4033 ± 0.0120** |
| jejunum | epithelial | DNA only | 0.4880 ± 0.0113 | 0.5761 ± 0.0102 | 0.0212 ± 0.0009 | 0.2239 ± 0.0094 |
| | | ATAC only | 0.5302 ± 0.0126 | 0.6798 ± 0.0055 | 0.0208 ± 0.0012 | 0.2369 ± 0.0104 |
| | | **DNA+ATAC** | **0.5939 ± 0.0105** | **0.7156 ± 0.0058** | **0.0180 ± 0.0007** | **0.3406 ± 0.0108** |
| | stromal (immune) | DNA only | 0.4157 ± 0.0153 | 0.5444 ± 0.0139 | 0.0100 ± 0.0006 | 0.1533 ± 0.0102 |
| | | ATAC only | 0.4706 ± 0.0073 | 0.6592 ± 0.0035 | 0.0095 ± 0.0007 | 0.1948 ± 0.0061 |
| | | **DNA+ATAC** | **0.5219 ± 0.0107** | **0.6895 ± 0.0053** | **0.0087 ± 0.0006** | **0.2624 ± 0.0079** |
| | stromal (other) | DNA only | 0.4679 ± 0.0131 | 0.5575 ± 0.0125 | 0.0136 ± 0.0005 | 0.1866 ± 0.0134 |
| | | ATAC only | 0.4587 ± 0.0041 | 0.6241 ± 0.0045 | 0.0137 ± 0.0008 | 0.1807 ± 0.0067 |
| | | **DNA+ATAC** | **0.5342 ± 0.0074** | **0.6671 ± 0.0065** | **0.0121 ± 0.0006** | **0.2746 ± 0.0083** |
| | ISC | DNA only | 0.4576 ± 0.0236 | 0.5701 ± 0.0150 | 0.0145 ± 0.0005 | 0.1922 ± 0.0193 |
| | | ATAC only | 0.5076 ± 0.0092 | 0.6604 ± 0.0053 | 0.0139 ± 0.0007 | 0.2273 ± 0.0116 |
| | | **DNA+ATAC** | **0.5589 ± 0.0158** | **0.6979 ± 0.0092** | **0.0125 ± 0.0005** | **0.3045 ± 0.0143** |

### Supplementary Figures

S1 10x Genomics multiome datasets . . . . . . . . . . . . . . . . . . . . . . . 11

S2 Model architecture . . . . . . . . . . . . . . . . . . . . . . . . . . . . . . . 12

S3 Naïve predictor performance on the PBMC dataset . . . . . . . . . . . . . . 12

S4 Cross-cell type performance within the PBMC dataset . . . . . . . . . . . . 13

S5 Cross-cell type performance within the brain dataset . . . . . . . . . . . . . 14

S6 Cross-cell type performance within the jejunum dataset . . . . . . . . . . . . 15

S7 Cross-cell type performance summary, evaluated on highly variable genes . . 16

S8 mRNA half-life and proximal pausing analysis in CD4 T cells . . . . . . . . . 16

S9 Comparison of PBMC GEx between multiome and single-cell datasets . . . . 17

S10 Cross-cell type performance within the PBMC single-cell dataset . . . . . . . 18

S11 Comparison of mean positional SHAP value correlation with chromatin accessibility between multiome and single-cell CD4 T cell models by channel . . . 19

S12 Mean log UMIs per cell by cell type in each dataset used in this study . . . . 20

S13 Relationship between dataset statistics and model performance . . . . . . . . 21

S14 Comparison of top 6-mers by mean attribution in the PBMC (sc) dataset . . 22

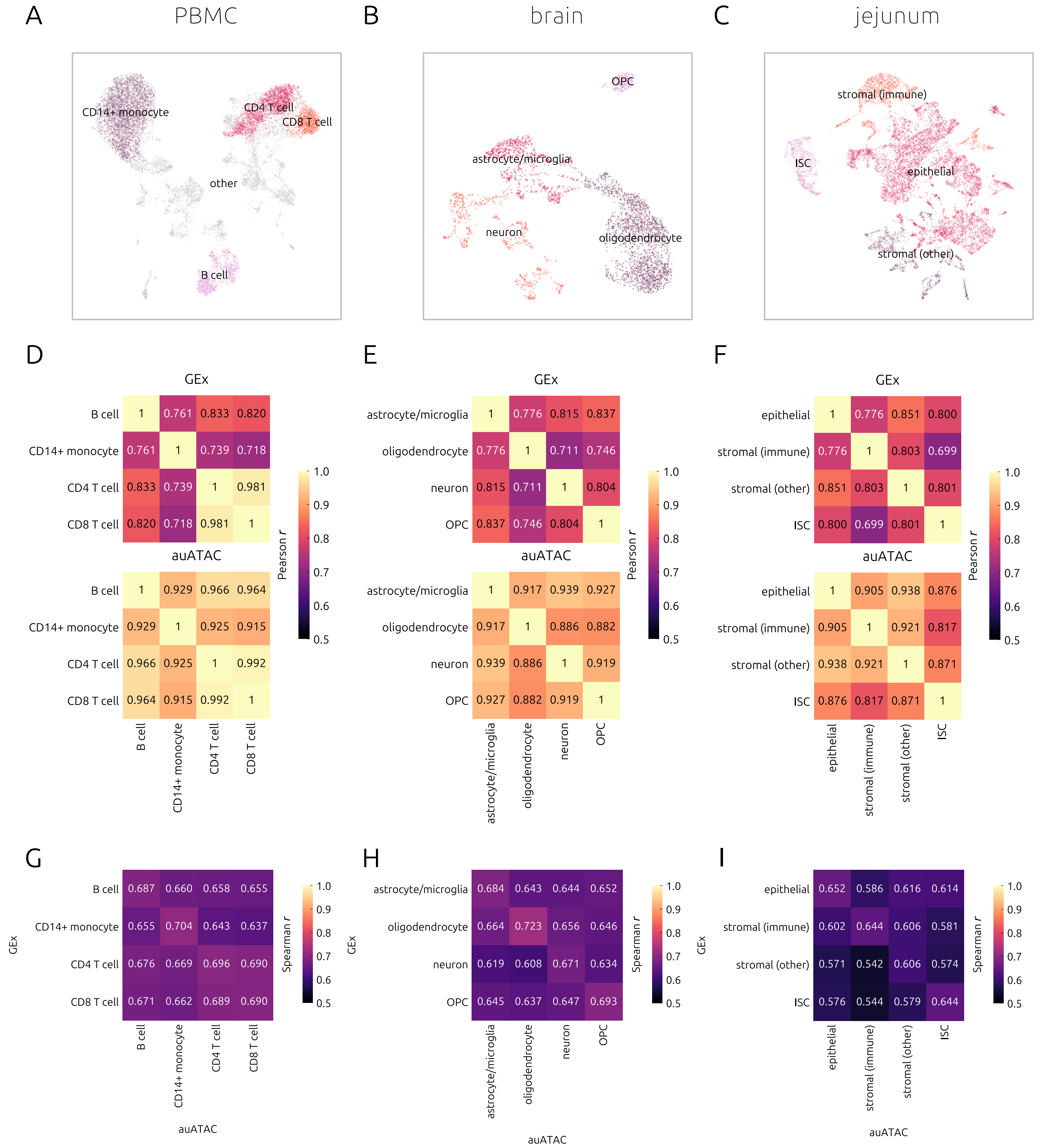

**Supplementary Figure S1:** 10x Genomics multiome datasets: PBMC (left), brain (center), and jejunum (right). A-C. Uniform manifold approximation and projection graphs of gene expression in 10x Genomics multiome datasets. The 4 major cell types used in each dataset are indicated in colour. D-F. Pearson correlation of GEx (top) and auATAC (bottom) between each major cell type in each multiome dataset. G-I. Spearman correlation between GEx and auATAC in each multiome dataset, within and between each major cell type.

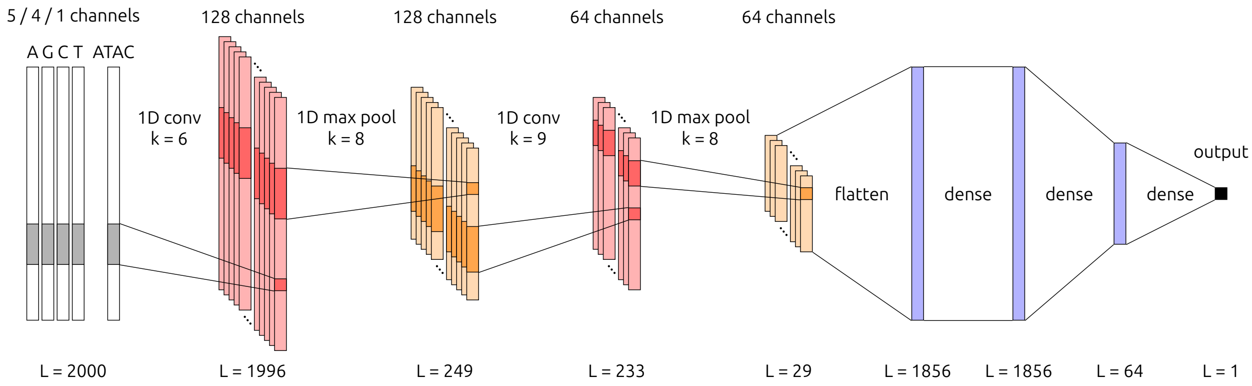

**Supplementary Figure S2:** Model architecture, adapted from Xpresso (Agarwal et al. 2020). Input channels may consist of 5, 4, or 1 channels, depending on the selected combination of DNA sequence and/or ATAC-seq track inputs. The first part of the model consists of 2 convolutional blocks in which each 1D convolution layer is followed by a 1D max pooling layer. Channel numbers and kernel sizes (k) are indicated for each layer. The second part of the model consists of 3 dense layers with ReLU activation, yielding a single output value.

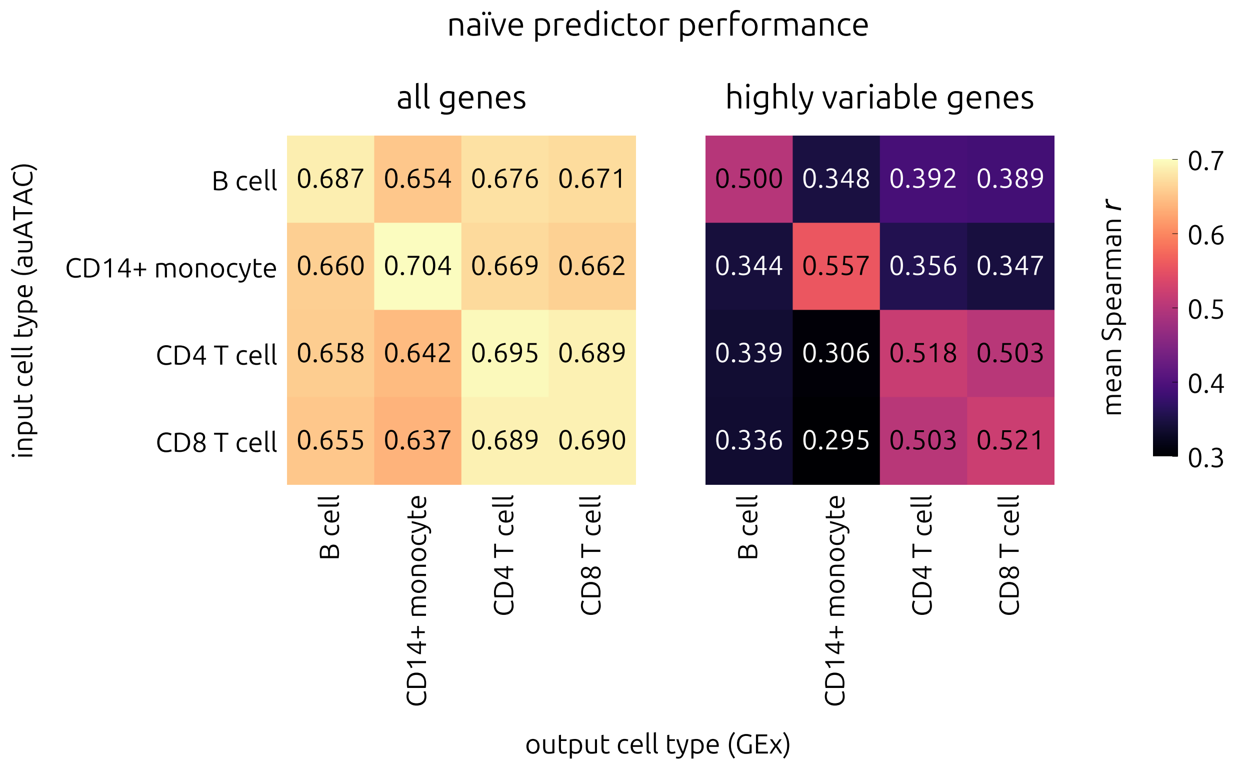

**Supplementary Figure S3:** Naïve predictor performance on the PBMC dataset. For each fold of 5-fold CV, the naïve predictor outputs the area under the input ATAC input track as a prediction for GEx. Mean Spearman correlation coefficients are shown for each combination of input and output cell type. On the left, evaluation results are shown using all genes in the test sets. On the right side, evaluation was restricted to only highly variable genes in the test sets.

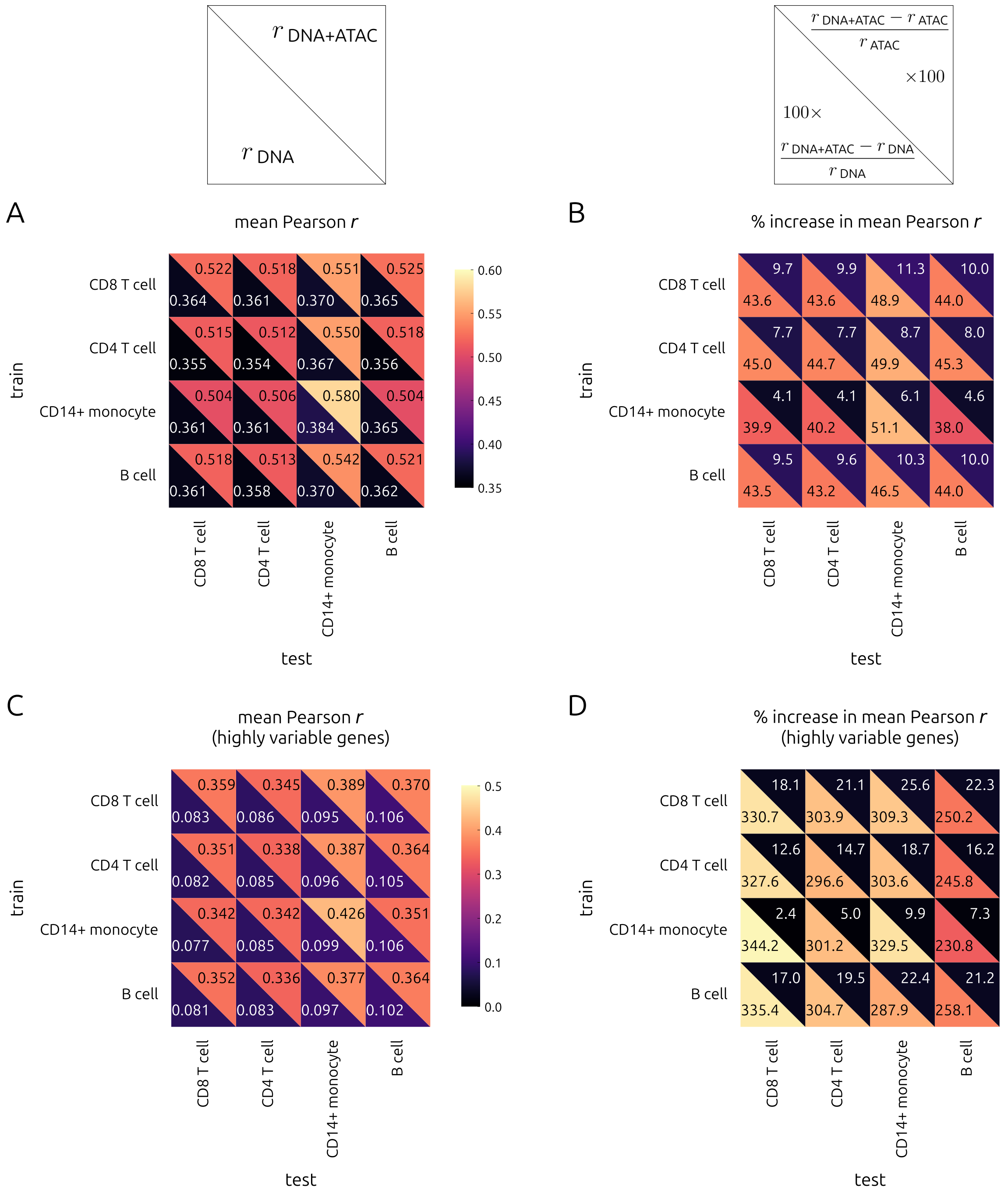

**Supplementary Figure S4:** Cross-cell type performance within the PBMC dataset. A. Mean Pearson $r$ of models trained on one cell type (rows) and tested on another (columns) for the PBMC dataset. Metrics for the DNA+ATAC model are shown in the top right triangle of each grid square. Metrics for the DNA-only model are shown in the bottom left triangle of each grid square. B. Same as A, but showing mean percent increase in Pearson $r$ of the DNA+ATAC model relative to DNA only (bottom left) and ATAC only (top right). C. Same as A, but evaluated on highly variable genes. D. Same as B, but evaluated on highly variable genes.

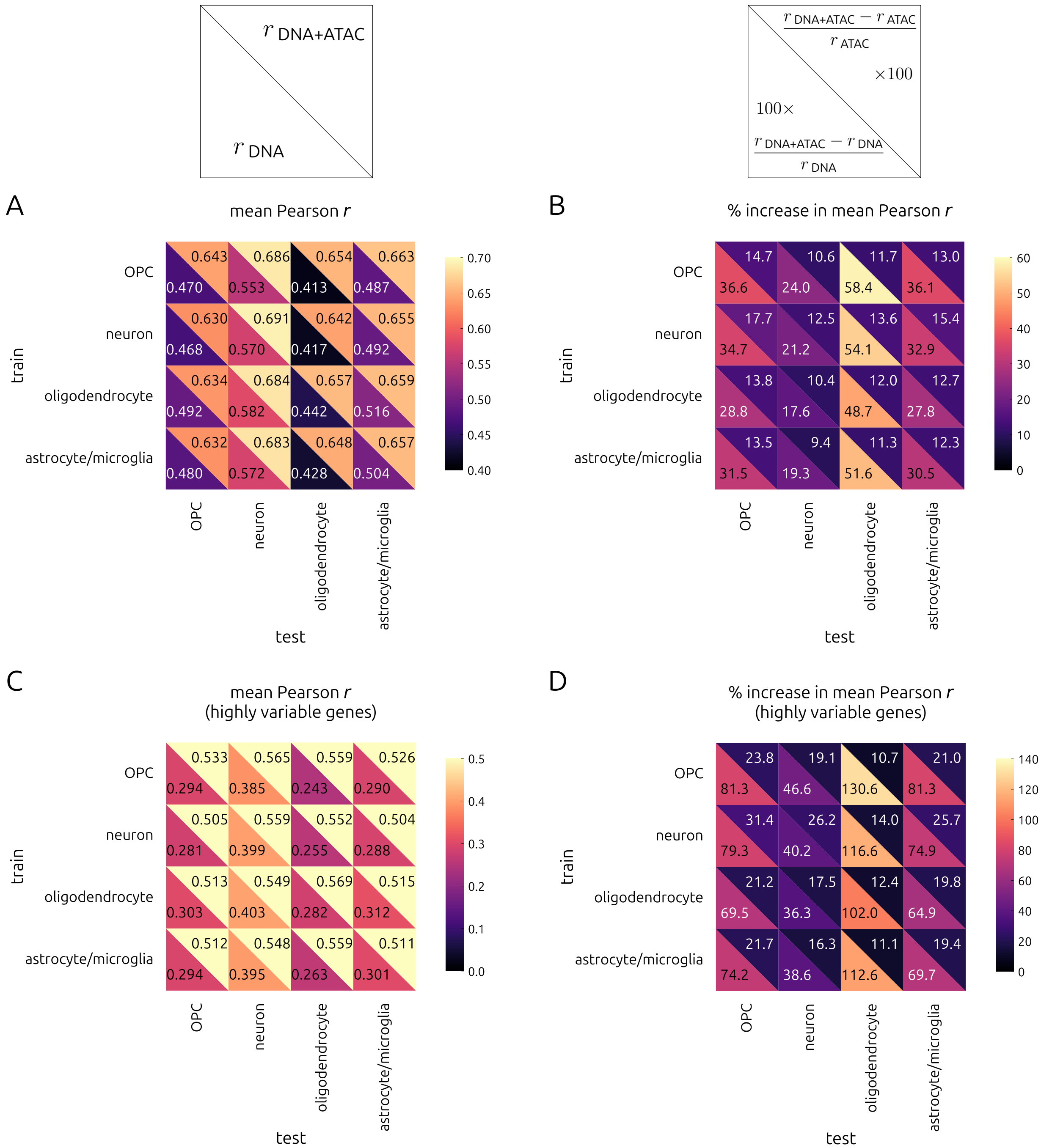

**Supplementary Figure S5:** Cross-cell type performance within the brain dataset. A. Mean Pearson $r$ of models trained on one cell type (rows) and tested on another (columns) for the brain dataset. Metrics for the DNA+ATAC model are shown in the top right triangle of each grid square. Metrics for the DNA-only model are shown in the bottom left triangle of each grid square. B. Same as A, but showing mean percent increase in Pearson $r$ of the DNA+ATAC model relative to DNA only (bottom left) and ATAC only (top right). C. Same as A, but evaluated on highly variable genes. D. Same as B, but evaluated on highly variable genes.

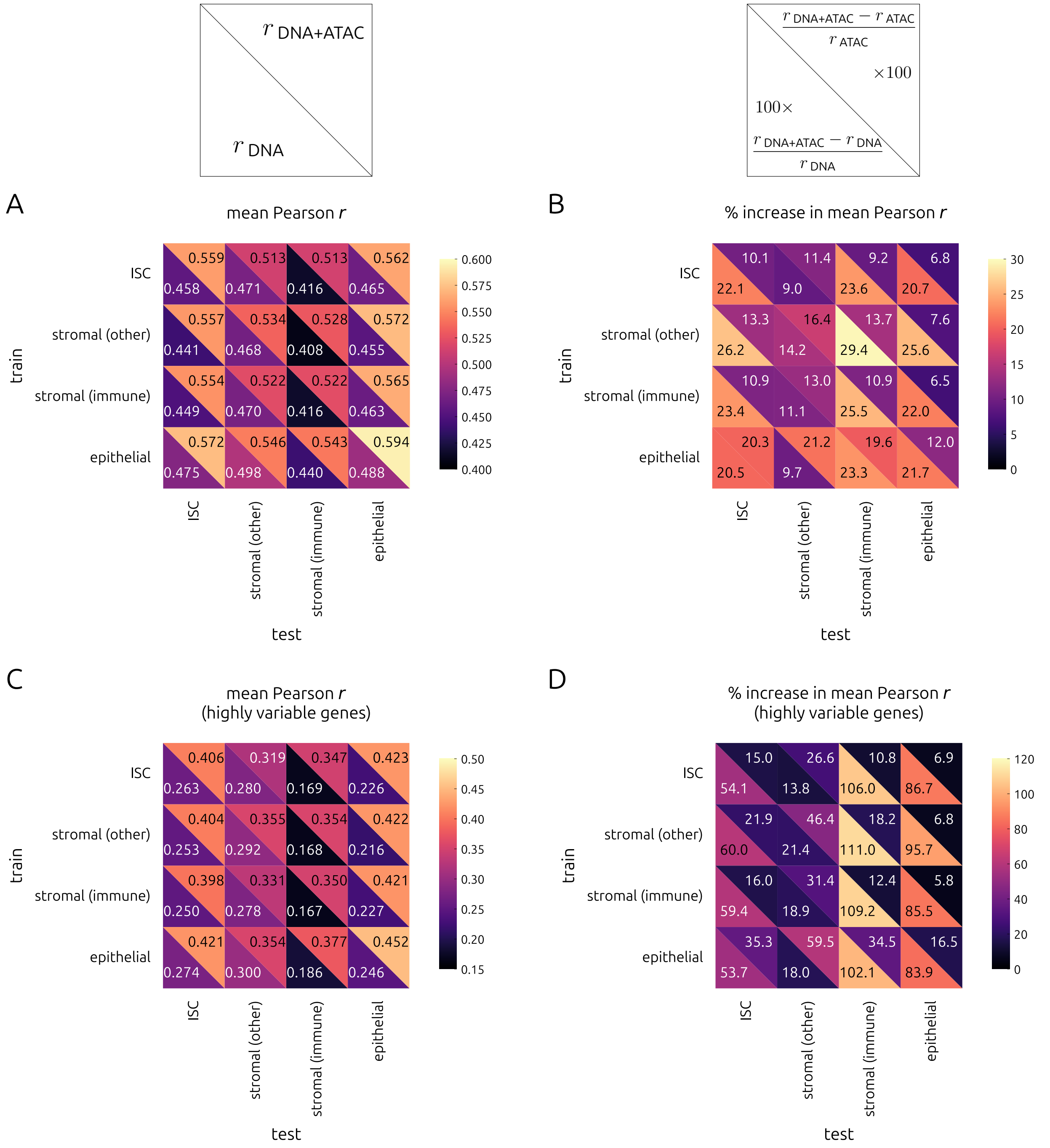

**Supplementary Figure S6:** Cross-cell type performance within the jejunum dataset. A. Mean Pearson $r$ of models trained on one cell type (rows) and tested on another (columns) for the jejunum dataset. Metrics for the DNA+ATAC model are shown in the top right triangle of each grid square. Metrics for the DNA-only model are shown in the bottom left triangle of each grid square. B. Same as A, but showing mean percent increase in Pearson $r$ of the DNA+ATAC model relative to DNA only (bottom left) and ATAC only (top right). C. Same as A, but evaluated on highly variable genes. D. Same as B, but evaluated on highly variable genes.

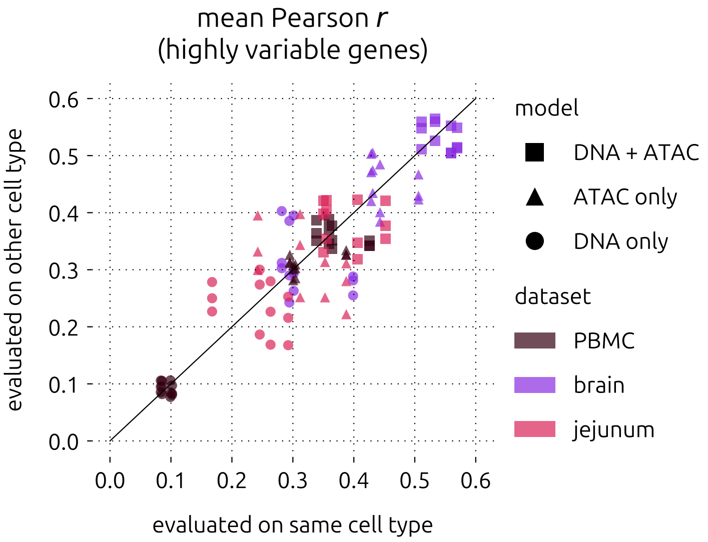

**Supplementary Figure S7:** Cross-cell type performance summary, evaluated on highly variable genes. Mean Pearson $r$ of models evaluated on held-out test sequences using another cell type from the same dataset (tissue) versus on the cell type used for training. Datasets are colour-coded. Marker shapes correspond to model types: DNA+ATAC (square), ATAC only (triangle), and DNA only (circle).

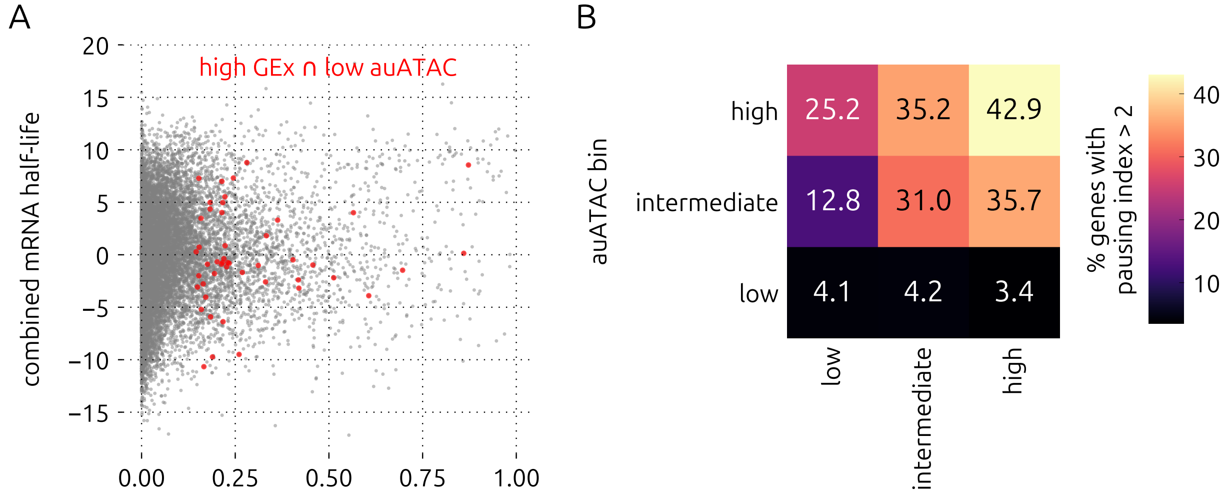

**Supplementary Figure S8:** mRNA half-life and proximal pausing analysis in CD4 T cells. A. Combined general mRNA half-life "meta-feature" [Agarwal and Kelley, 2022] versus GEx in CD4 T cells from the multiome PBMC dataset. Genes in the high GEx and low auATAC category are coloured red. B. The percentage of genes in each CD4 T cell category with a pausing index greater than 2, as determined from PRO-seq data of human PBMC CD4 T cells [Danko et al., 2018].

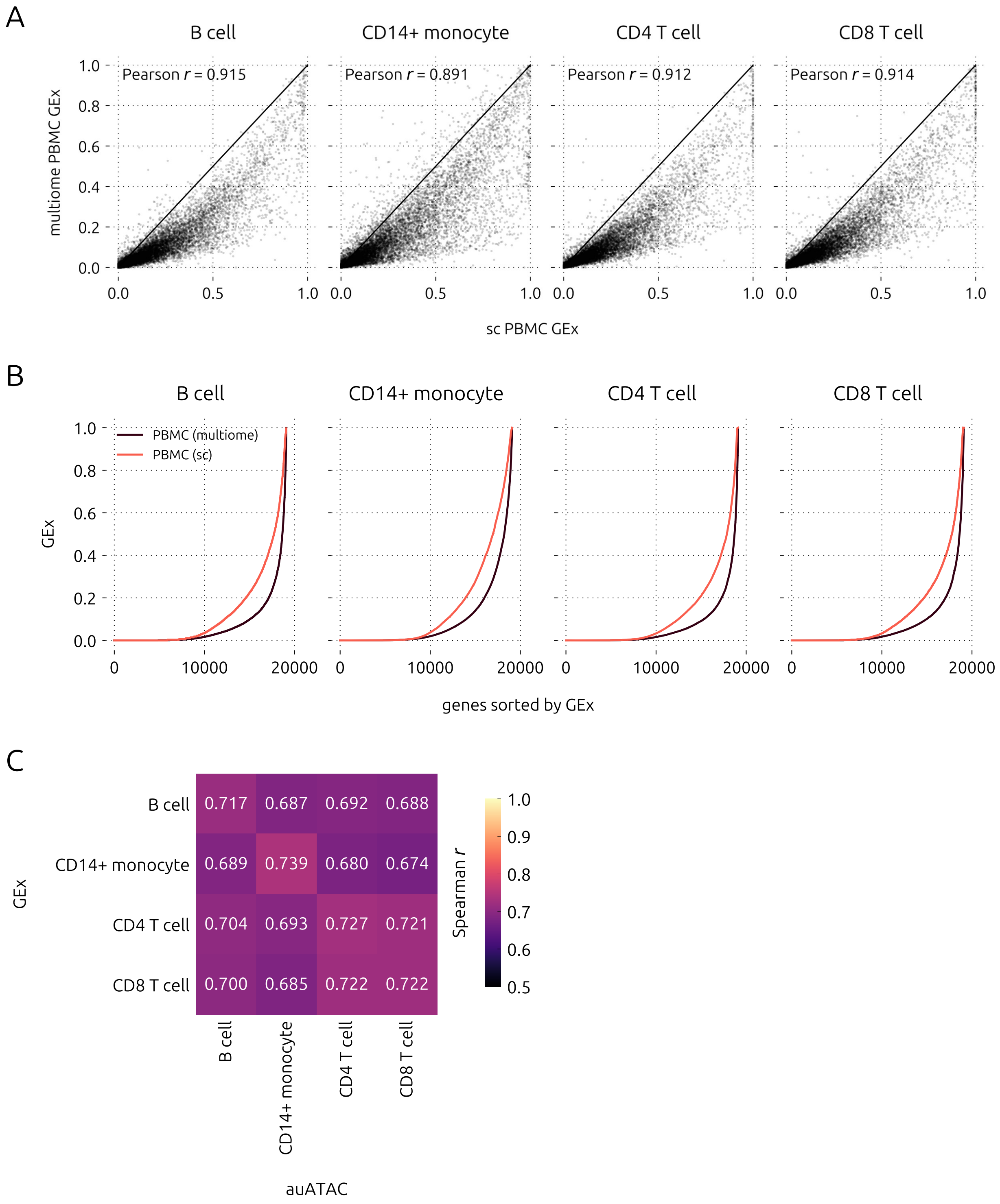

**Supplementary Figure S9:** Comparison of PBMC GEx between multiome and single-cell datasets. A. Multiome versus single-cell GEx in each cell type. B. Ranked GEx values in multiome and single-cell datasets. C. Spearman correlation between sc GEx and multiome auATAC for each major cell type.

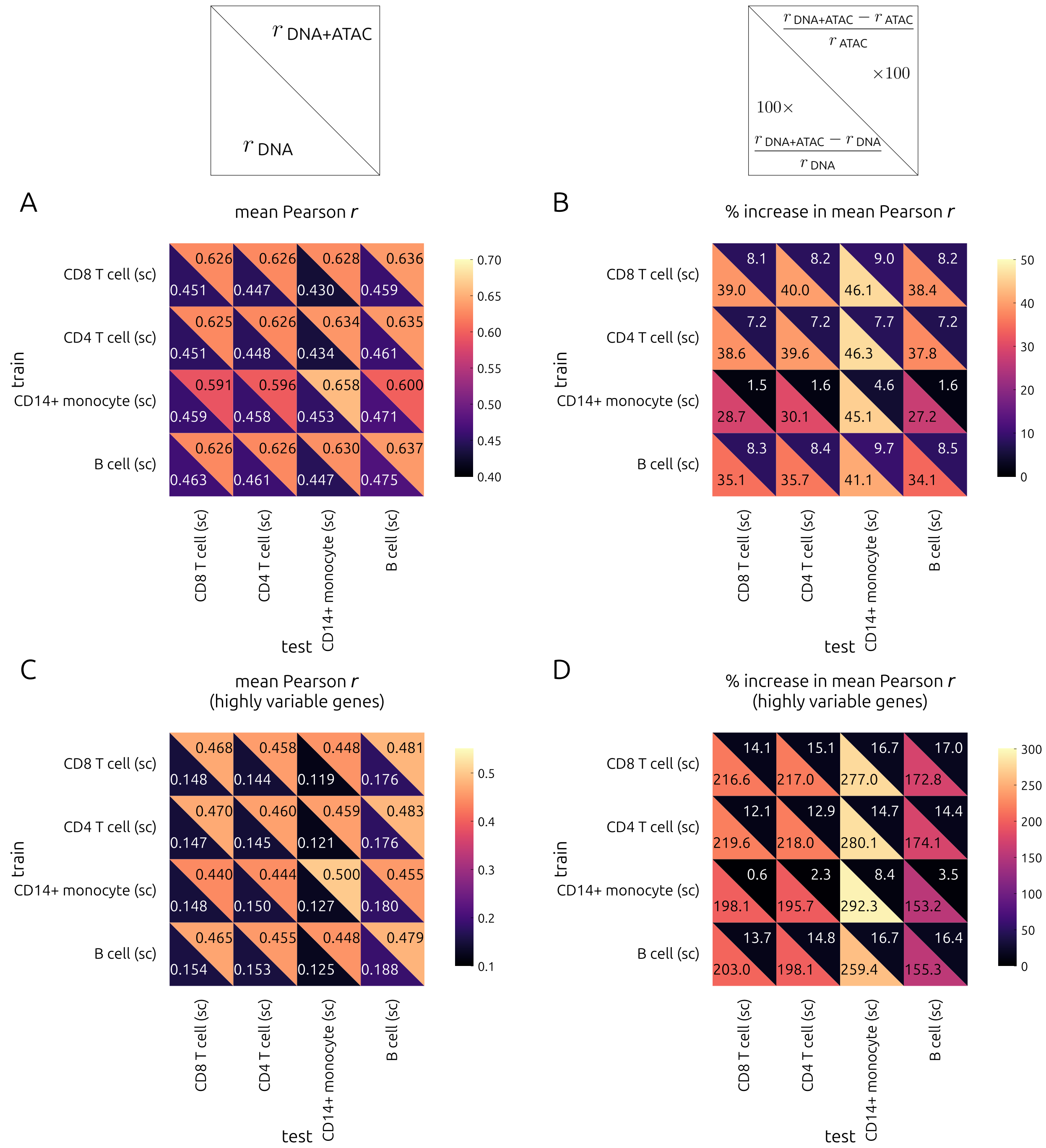

**Supplementary Figure S10:** Cross-cell type performance within the PBMC single-cell dataset, using ATAC data from the PBMC multiome dataset. A. Mean Pearson $r$ of models trained on one cell type (rows) and tested on another (columns) for the PBMC single-cell dataset. Metrics for the DNA+ATAC model are shown in the top right triangle of each grid square. Metrics for the DNA-only model are shown in the bottom left triangle of each grid square. B. Same as A, but showing mean percent increase in Pearson $r$ of the DNA+ATAC model relative to DNA only (bottom left) and ATAC only (top right). C. Same as A, but evaluated on highly variable genes. D. Same as B, but evaluated on highly variable genes.

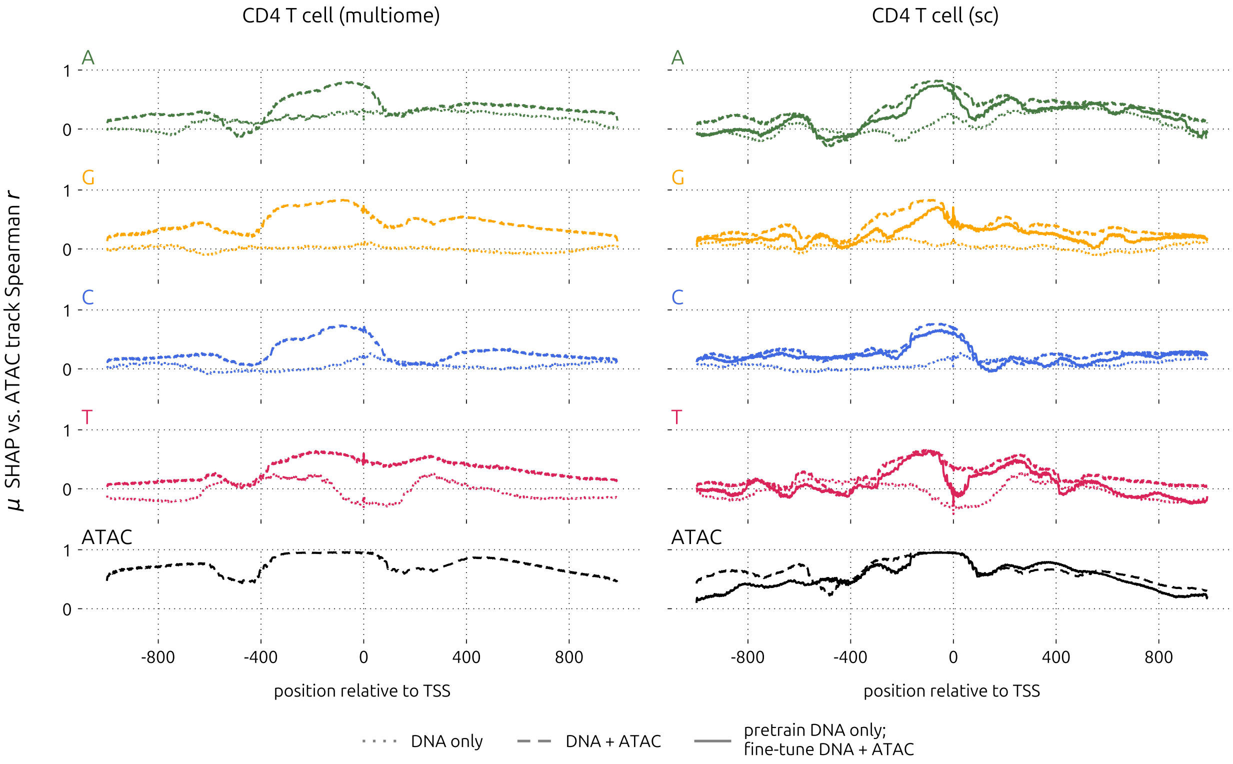

**Supplementary Figure S11:** Comparison of mean positional SHAP value correlation with chromatin accessibility between multiome (left) and single-cell (right) CD4 T cell models by channel. Dotted, dashed, and solid lines indicate DNA-only, DNA+ATAC, and DNA+ATAC with DNA-only pretraining, respectively.

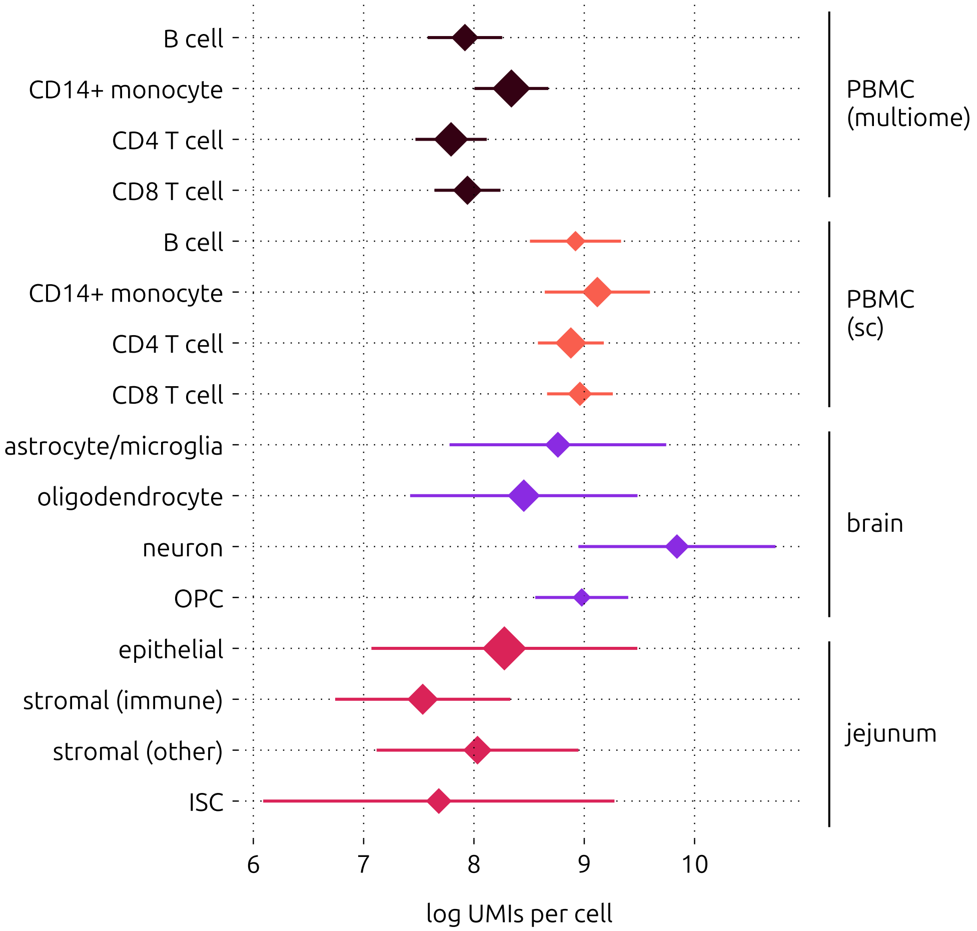

**Supplementary Figure S12:** Mean log UMIs per cell by cell type in each dataset used in this study. Marker area is proportional to the number of cells pooled for that cell type. Error bars represent standard deviation.

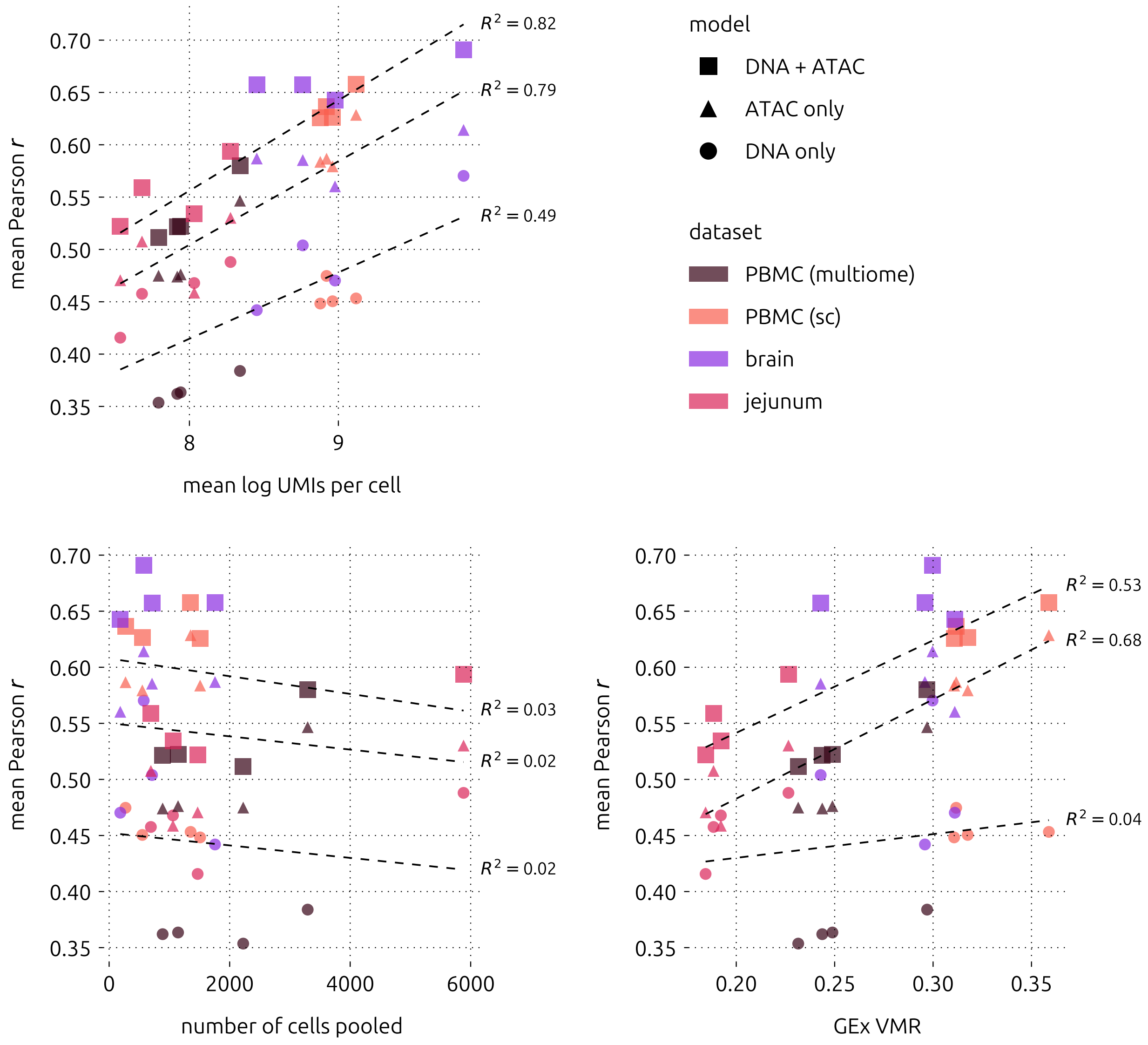

**Supplementary Figure S13:** Relationship between dataset statistics and model performance. Top: mean Pearson correlation versus mean log UMIs per cell. Bottom left: mean Pearson correlation versus number of cells pooled. Bottom right: mean Pearson correlation versus variance-to-mean ratio (VMR) of GEx in that dataset. Datasets are colour-coded. Marker shapes correspond to model types: DNA+ATAC (square), ATAC only (triangle), and DNA only (circle). For each model type, the line of best fit and its coefficient of determination ($R^2$) is shown.

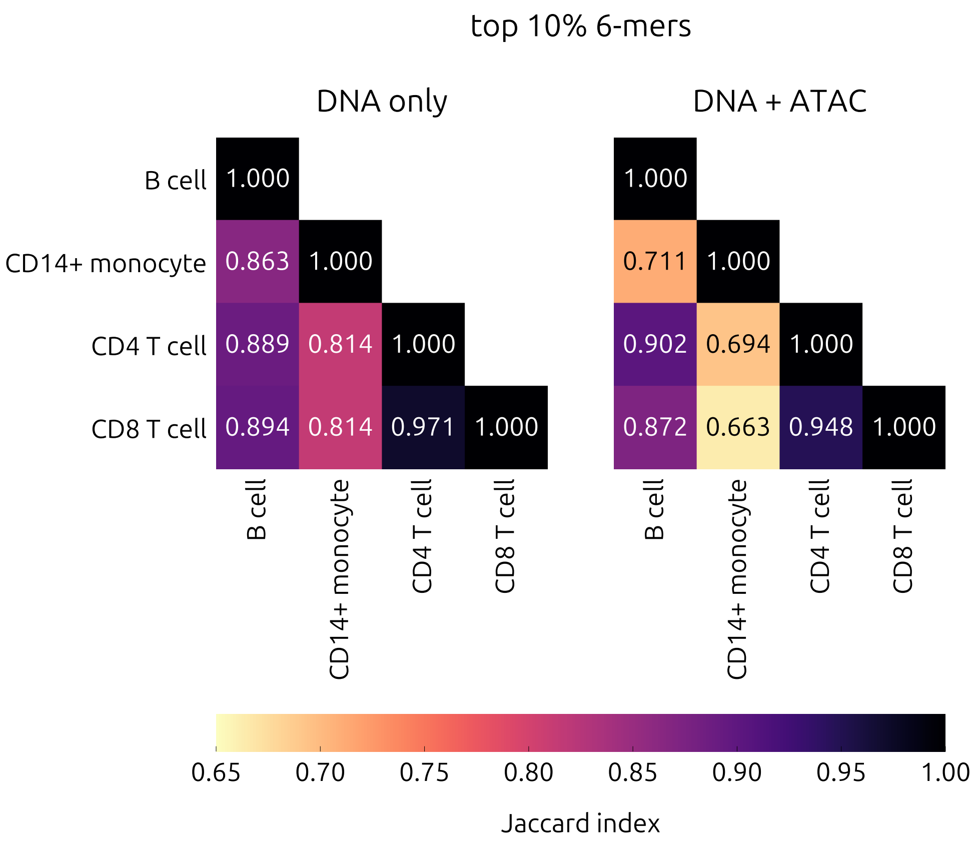

**Supplementary Figure S14:** Comparison of top 6-mers by mean attribution in the PBMC (sc) dataset, using the Jaccard index.
